# Comparative sensitivity of *in vitro* acute and chronic apical and transcriptomic points of departure for perfluorooctanoic acid in human vascular endothelial cells

**DOI:** 10.1101/2024.12.24.628084

**Authors:** Marija Opacic, Bojana Stanic, Nebojsa Andric

## Abstract

*In vitro* human cell models and new approach methodologies offer valuable alternatives to animal testing, providing human-relevant data for chemical risk assessment. However, the impact of exposure duration and biological levels on the sensitivity of responses in human cells *in vitro* remains underexplored. In this study, we employed benchmark concentration modeling to derive and compare points of departure (PODs) for apical (A-POD) and transcriptomic (T-POD) changes in human vascular endothelial cells EA.hy926 following short-term (48 h) and long-term (6 and 12 weeks) exposure to 1, 10, and 100 µM and 1, 10, and 100 nM perfluorooctanoic acid (PFOA), respectively. We found that A-POD could not be calculated for short-term exposure; however, A-PODs after 6 and 12 weeks were determined to be 3.7 nM and 5.4 nM, respectively. mRNA sequencing revealed a significant number of differentially expressed genes in PFOA-exposed EA.hy926 cells across all time points. T-POD values after 6 and 12 weeks (4.1 nM and 22.1 nM, respectively) demonstrated greater sensitivity than the acute T-POD (6.3 µM) and were comparable to the chronic A-POD. Functional gene analysis revealed that transcription was a sensitive, yet general molecular pathway affected by acute exposure, while the IL-17 pathway and extracellular matrix and cytokine-cytokine interactions were implicated after 6 and 12 weeks of exposure to PFOA, respectively. In conclusion, chronic PODs proved to be more informative than acute PODs for chemical risk assessment in PFOA-exposed human endothelial cells. Furthermore, transcriptomics data from long-term exposure can elucidate early molecular pathways associated with chemical exposure, highlighting their potential to connect toxicity effects across different biological levels.

## 1. Introduction

Perfluorooctanoic acid (PFOA; IUPAC name: 2,2,3,3,4,4,5,5,6,6,7,7,8,8,8-pentadecafluorooctanoic acid, CAS# 335-67-1) is a synthetic chemical belonging to the class of per- and polyfluoroalkyl substances (PFAS). PFAS, including PFOA, have been widely synthesized and used in various industries and consumer products since the 1940s, but have recently gained attention as significant environmental contaminants (Wee and Aris, 2023). In recent years, PFOA and other notable PFAS, such as perfluorooctanesulfonic acid (PFOS), have been phased out and replaced with alternative chemicals by major manufacturing companies (Land et al., 2018). Despite these efforts, because of their extreme persistence (often referred to as “forever chemicals”) they continue to be found in both the environment and the human body, posing substantial risks to human and environmental health. Exposure to PFOA occurs through contaminated water, soil, air, and food, affecting humans, livestock, and wildlife alike. In the general population, reported blood plasma concentrations of PFOA range from 1 to 10 ng/mL (Hölzer et al., 2021). In a population exposed to PFAS-contaminated groundwater, the median serum concentration of PFOA was found to be 49 ng/mL, with an estimated half-life of approximately 2.36 years, shorter in women than in men (Batzella et al., 2024). Among adolescents and young adults exposed to high levels of PFAS through drinking water, PFOA exhibited the highest serum concentration, with a median value of 44.4 ng/mL (Pitter et al., 2020). A study of over 46,000 community residents aged 18 and older from the Mid-Ohio Valley, who had been exposed to various levels of PFOA through contaminated drinking water and food for decades, revealed a median blood PFOA level of 27 ng/mL, with concentrations ranging from as low as 0.25 ng/mL to nearly 18,000 ng/mL (Steenland et al., 2009).

PFOA is associated with a wide array of adverse health effects, including reproductive and developmental toxicity, hepatotoxicity, renal toxicity, immunotoxicity, neurotoxicity, genotoxicity, and carcinogenicity (Li et al., 2017). Research indicates that PFOA can negatively impact general growth and development, as well as induce sex-specific alterations in pubertal maturation (Li et al., 2017), and can also exert direct adverse effects on steroid hormone production and indirect effects on ovarian function, leading to classification of PFOA as a potential endocrine disruptor (White et al., 2011). However, little is known about the effects of PFOA on human vascular endothelial cells and the vascular system in general. The inner lining of arteries, veins, and capillaries is made up of over one trillion endothelial cells that are continuously exposed to various stimuli, including environmental chemicals (Krüger-Genge et al., 2019). Increasing evidence suggests that exposure to harmful substances in the blood can lead to endothelial cell damage and, consequently, endothelial dysfunction. Experimental studies have shown that vascular endothelial cells can respond adversely to environmental chemicals, such as bisphenol A and phthalates, potentially compromising vascular function (Kokai et al., 2024; Kokai et al., 2020; Kokai et al., 2022; Kong et al., 2022; Stanic et al., 2023; Stanic et al., 2024; Stanic et al., 2022). In contrast, studies specifically investigating the impact of PFOA on endothelial cells are limited. For instance, one study explored whether exposure to PFOA could induce morphological changes in the vascular 3D network structure (Poteser et al., 2020). Other studies have examined the effects of PFOA on the plasma kallikrein-kinin system, which regulates vascular permeability (Liu et al., 2018), and whether gestational exposure to PFOA could inhibit early placenta development through the shrinkage of labyrinth vessels (Jiang et al., 2020).

It is clear that the effects of PFOA on vascular endothelial cells, as well as the sensitivity of these responses, have not been sufficiently studied to date. This gap is critical because regulatory agencies worldwide are developing guidelines for PFAS and need to identify exposure levels that may pose risks to human health. Establishing these levels is essential for setting safety limits to protect public health and reduce the social and economic burdens associated with chemical exposure. An essential aspect of this evaluation is the use of sensitive, precise, and efficient testing methodologies. Currently, assessment largely relies on human biomonitoring and epidemiological studies, alongside observations of adverse effects in animal studies. However, these approaches have several limitations, making the evaluation of chemical adverse effects less efficient. Comprehensive datasets obtained through various approaches – such as long-term exposure studies, consideration of gender differences, the use of different animal species, and testing for low-dose effects – are necessary for developing robust regulatory guidelines for PFOA. One of the approaches that can facilitate chemical risk characterization is the identification of the point of departure (POD). Traditionally, the POD is derived from no-observed-adverse-effect level (NOAEL) or lowest-observed-adverse-effect level (LOAEL) analyses, which estimate the quantity of a substance above which adverse effects are expected to occur in humans (Izadi et al., 2012). More recently, benchmark dose (BMD) analysis has replaced the NOAEL/LOAEL approach as the preferred method for identifying PODs in chemical risk assessment (Johnson et al., 2014). The BMD is calculated using dose-response curve, taking into account the shape of the curve to establish a lower confidence limit of the BMD (BMDL), which corresponds to a given increase in response over the background level (Guo and Mei, 2018). The BMDL thus serves as the POD. To derive a Maximum Contaminant Level Goal for PFOA and PFOS in drinking water, the United States Environmental Protection Agency (US EPA) employed various approaches. These included identifying critical toxicity endpoints of PFOA and PFOS exposure, i.e., POD, utilizing BMD modeling to determine an internal POD, employing pharmacokinetic modeling to establish a human equivalent dose, and applying an interspecies uncertainty factor to account for variability in cellular response and pharmacokinetics of PFAS across the human population. As a result, interim drinking water health advisories for PFOS and PFOA were significantly lowered, from 70 parts per trillion (ppt) to 0.04-0.2 ppt (Perez et al., 2023). Innovative approaches, such as BMD modeling, can help identify thresholds of molecular change. However, several critical questions regarding chemical impact assessment remain unresolved, including the choice of model organism, duration of exposure, and the level of biological organization – specifically, the use of toxicogenomic outputs versus traditional endpoint measurements, which are still regarded as the gold standard in toxicology testing. While some of these questions have been addressed in *in vivo* studies using animal models (Bhat et al., 2013; Bianchi et al., 2021; Gwinn et al., 2020; Johnson et al., 2020; Thomas et al., 2013), significant gaps persist in the application of *in vitro* human cell models.

The aim of this study was to utilize the BMD approach to compare comprehensive datasets obtained from HTS toxicogenomic analysis with traditional toxicological endpoints in *in vitro* human cells model. We achieved this objective through various methodological approaches, including both short-term, acute (48 h) and long-term, chronic (6 and 12 weeks) *in vitro* exposure of human vascular endothelial cells to higher concentrations of PFOA (1 µM, 10 µM, and 100 µM) and real-life human exposure concentrations (1 nM, 10 nM, and 100 nM), respectively. In both exposure scenarios, we collected traditional toxicological apical data alongside toxicogenomics data, which we subsequently analyzed using BMD modeling. Our study not only provides novel insights into the effects of PFOA on human vascular endothelial cells, contributing to the growing understanding of PFOA-induced vascular toxicity, but also enhances the understanding of how different *in vitro* approaches can be integrated into chemical risk assessment.

## 2. Material and methods

### 2.1 Chemicals

PFOA (chemical purity ≥95%), Dulbecco’s Modified Eagle’s Medium (DMEM) containing 4 mM L-glutamine and 4.5 g/L glucose, RPMI-1640 medium with 2 mM L-glutamine, 4.5 g/L glucose, and 25 mM HEPES, Hanks’ Balanced Salt Solution (HBSS) without phenol red, penicillin (10 000 IU/mL)-streptomycin (10 mg/mL) mixture, 0.25% trypsin-ethylenediaminetetraacetic acid (EDTA) solution, bovine serum albumin (BSA), dimethyl sulfoxide (DMSO), sulforhodamine B (SRB), trichloroacetic acid, fibronectin from bovine plasma (1 mg/mL in 0.05 M Tris-buffered saline, pH 7.5), Triton X-100, and Trypan Blue solution (0.4%) were obtained from Sigma-Aldrich Chemie GmbH (Steinheim, Germany). Fetal bovine serum (FBS), HEPES, HAT supplement (5 mM sodium hypoxanthine, 20 µM aminopterin, 0.8 mM thymidine), calcein AM, fluorescein isothiocyanate (FITC)-dextran (FD40, Mw 40 kDa), 2′,7′-dichlorodihydrofluorescein diacetate (H_2_DCFDA), propidium iodide (chemical purity ≥95%), RNAse A, TRIzol Reagent, and alamarBlue™ Cell Viability Reagent were purchased from Thermo Fisher Scientific (Waltham, MA, USA). Annexin V-FITC was from Elabscience (Houston, TX, USA). Cultrex Reduced Growth Factor Basement Membrane Extract (BME), PathClear, was obtained from R&D Systems/Bio-Techne (Minneapolis, MN, USA). VECTASHIELD^®^ PLUS Antifade Mounting Medium with DAPI was from Vector Laboratories (Newark, CA, USA). All other chemicals were of analytical grade.

### 2.2 Cell culture

The human macrovascular endothelial cell line EA.hy926 was maintained as a monolayer culture at 37 °C in a humidified atmosphere containing 5% CO_2_, as previously described (Kokai et al., 2020). The complete culture medium consisted of DMEM supplemented with 10% FBS, 1% penicillin-streptomycin solution, 1.5 g/L NaHCO_3_, 0.11 g/L sodium pyruvate, 10 mM HEPES buffer, and 4% HAT. The medium was refreshed every three to four days, with subculturing performed when the cells reached ∼90% confluency. Cells were washed twice with phosphate-buffered saline (PBS) and then trypsinized using 0.25% trypsin-EDTA for 4 min. Following this, complete culture medium was added, and the cells were centrifuged at 400 × g for 5 minutes. The resulting suspension was mixed with Trypan blue solution and counted using an automated cell counter.

The human pro-monocytic cell line U937 was grown as a suspension culture at 37 °C in a humidified atmosphere with 5% CO_2_, as detailed in the Supplementary Material of our previous publication (Stanic et al., 2022). Cell suspensions at appropriate densities were utilized for both cell line propagation (up to 0.3 × 10^6^ cells per mL) and for conducting the monocyte-endothelial adhesion assay following short-term (acute) and long-term (chronic) exposure of EA.hy926 cells to PFOA.

### 2.3 Short-term exposure of EA.hy926 cells to PFOA

EA.hy926 cells were cultured in 75 cm^2^ flasks in complete culture medium under standard conditions. Cells were subcultured twice weekly – 1.8 × 10^6^ cells were returned to the flasks on Mondays, and 2.8 × 10^6^ cells on Fridays. For short-term exposure, EA.hy926 cells were plated in appropriate culture plates or dishes, and 24 h after seeding exposed to either vehicle (0.05% DMSO – control group) or three micromolar concentrations of PFOA (1 µM, 10 µM, and 100 µM, in 0.05% DMSO – treatment groups) for 48 h.

### 2.4 Long-term exposure of EA.hy926 cells to PFOA

Three different stock vials of EA.hy926 cells cryopreserved on different dates (biological replicates) were thawed and cultured in separate 25 cm^2^ flasks under standard conditions for two weeks. The cells from each flask were then divided into four flasks, resulting in a total of 12 flasks. Over the subsequent 12 weeks, EA.hy926 cells were subcultured twice a week – 0.95 × 10^6^ cells per flask were replated on Tuesdays, and 0.75 × 10^6^ cells on Fridays. Three h after seeding, cells were exposed to either vehicle (0.05% DMSO – control group) or three nanomolar concentrations of PFOA (1 nM, 10 nM, and 100 nM, in 0.05% DMSO – treatment groups) (Supplementary Figure 1). As specified in certain experiments, treatments were also continued in cell culture plates and dishes.

### 2.5 Apical data

The following apical points were analyzed after the short-term and long-term exposure of EA.hy926 cells to PFOA: metabolic activity, apoptosis and necrosis, cell cycle progression, endothelial permeability, monocyte adhesion to the endothelial cell monolayer, cell adhesion to the extracellular matrix, cell migration, endothelial tube formation, and the production of reactive oxygen species (ROS). Detailed protocols for these assays are provided in the Supplementary Material.

### 2.6 Trancriptomics

For short-term exposure, EA.hy926 cells were plated in 35-mm dishes (0.6 × 10^6^ cells per dish) and exposed to either vehicle (0.05% DMSO) or three micromolar concentrations of PFOA (1 μM, 10 μM, and 100 μM PFOA) the following day. After 48 h of exposure, the cells were rinsed with PBS, collected in 0.5 mL of TRIzol, and the RNA was extracted afterward. In long-term exposure experiments, EA.hy926 cells originating from each flask after 6 and 12 weeks of exposure to either vehicle (0.05% DMSO) or three nanomolar concentrations of PFOA (1 nM, 10 nM, and 100 nM PFOA) were collected directly during subculturing into clean RNase-, DNase-, and pyrogen-free 1.5 mL microcentrifuge tubes (∼1 × 10^6^ cells cells per tube). Cells were centrifuged at 400 × g for 5 minutes, rinsed with PBS, centrifuged again, and stored in 0.5 mL of TRIzol at –80 °C for subsequent RNA isolation. Three replicates of control and each treatment group per experiment were submitted to BGI Europe (Poland) for mRNA sequencing (RNA-Seq) and further analysis. RNA quantity and integrity were assessed using an Aglient 4150 Bioanalyzer (Agilent Technologies, Santa Clara, CA, USA). All samples had an RNA integrity number greater than 9.5, indicating very high quality. RNA-Seq was conducted on the DNBSEQ platform, with sequencing length PE150 and an average yield of 6.65G data per sample. Sample reads were trimmed to remove any reads with more than 5% unknown base content, along with adapters and low-quality bases. HISAT was used to align the clean reads to the reference genome (species: *Homo sapiens*; source: NCBI; reference genome version: GRCh38.p14, https://www.ncbi.nlm.nih.gov/datasets/genome/GCF_000001405.40/, accessed 18 October 2024), while Bowtie2 was used to align the clean reads to reference genes. The average alignment ratio of the sample comparison genome was 98.74%, whereas the average alignment of the gene set was 85.92%; a total of 18048 genes were detected. An initial bioinformatics analysis was performed by BGI. Normalized gene counts were further used for benchmark concentration (BMC) modeling.

### 2.7 Benchmark concentration modeling

BMC modeling was conducted on two sets of data (probes): apical points and log2-transformed DESeq2 normalized gene counts using the BMDExpress software v3.20.0106 available at https://github.com/auerbachs/BMDExpress-3. BMDExpress efficiently analyzes large transcriptomics datasets and other data types to select probes with concentration-response patterns and calculate BMC values, including the BMC lower confidence limit (BMCL) and BMC upper confidence limit (BMCU). These values represent concentrations that induce a predetermined change in the response rate of an adverse effect. Probes were pre-filtered using the Williams Trend Test with 500 permutations, a linear 1.5-fold change, and a significance threshold of *p* <0.05 to select the probes with specific dose responses. Probes passing the Williams Trend Test were further modeled using the US EPA’s BMDS online tool (https://bmdsonline.epa.gov/) with the following parameters: benchmark response (BMR) type as standard deviation (SD) with a BMR factor of 1 SD, model selection for BMC, BMCL, and BMCU in the best model, and best polynomial model test using Nested Chi-Square, with a *p*-value cut-off of 0.05. The BMC data were further filtered based on the following criteria: BMC greater than the concentrations used in the analysis, and a BMCU to BMCL ratio greater than 40. Gene accumulation was created in BMDExpress3, while density plots were created using the R program.

### 2.8 Gene enrichment analysis

Functional analysis of genes with calculated BMCLs (PODs) was performed using DAVID Bioinformatics Resources 6.8 (https://david.ncifcrf.gov/home.jsp). Gene enrichment analysis was conducted in DAVID using the default settings for *Homo sapiens*, with enrichment significance determined by the false discovery rate. Functional modules with at least five hits were represented using the ggplot2 package in R.

## 3. RESLUTS

### 3.1 Apical data

#### 3.1.1 Apoptosis and necrosis

Both short-term and long-term exposure of EA.hy926 cells to PFOA did not affect metabolic activity across any of the tested concentrations (Supplementary Figure 2). However, a 12-week exposure to 100 nM PFOA resulted in approximately a 17% reduction in cell proliferation (data not shown). To further investigate the effects of PFOA, we examined whether short-term and long-term exposures to selected concentrations induced apoptosis or necrosis in EA.hy926 cells. As illustrated in Figure 1, short-term exposure to 1 μM, 10 μM, and 100 μM PFOA did not trigger either apoptosis or necrosis. On the other hand, we noted a decline in live cell counts after 6 weeks of exposure to 1 nM PFOA and after 12 weeks of exposure to 1 nM, 10 nM, and 100 nM PFOA. The number of apoptotic cells increased only after 12 weeks of exposure to 1 nM and 100 nM PFOA. Additionally, necrotic cell counts rose after 6 weeks of exposure to 1 nM PFOA and after 12 weeks at all three concentrations of PFOA (Figure 1A and B).

**Figure 1.**
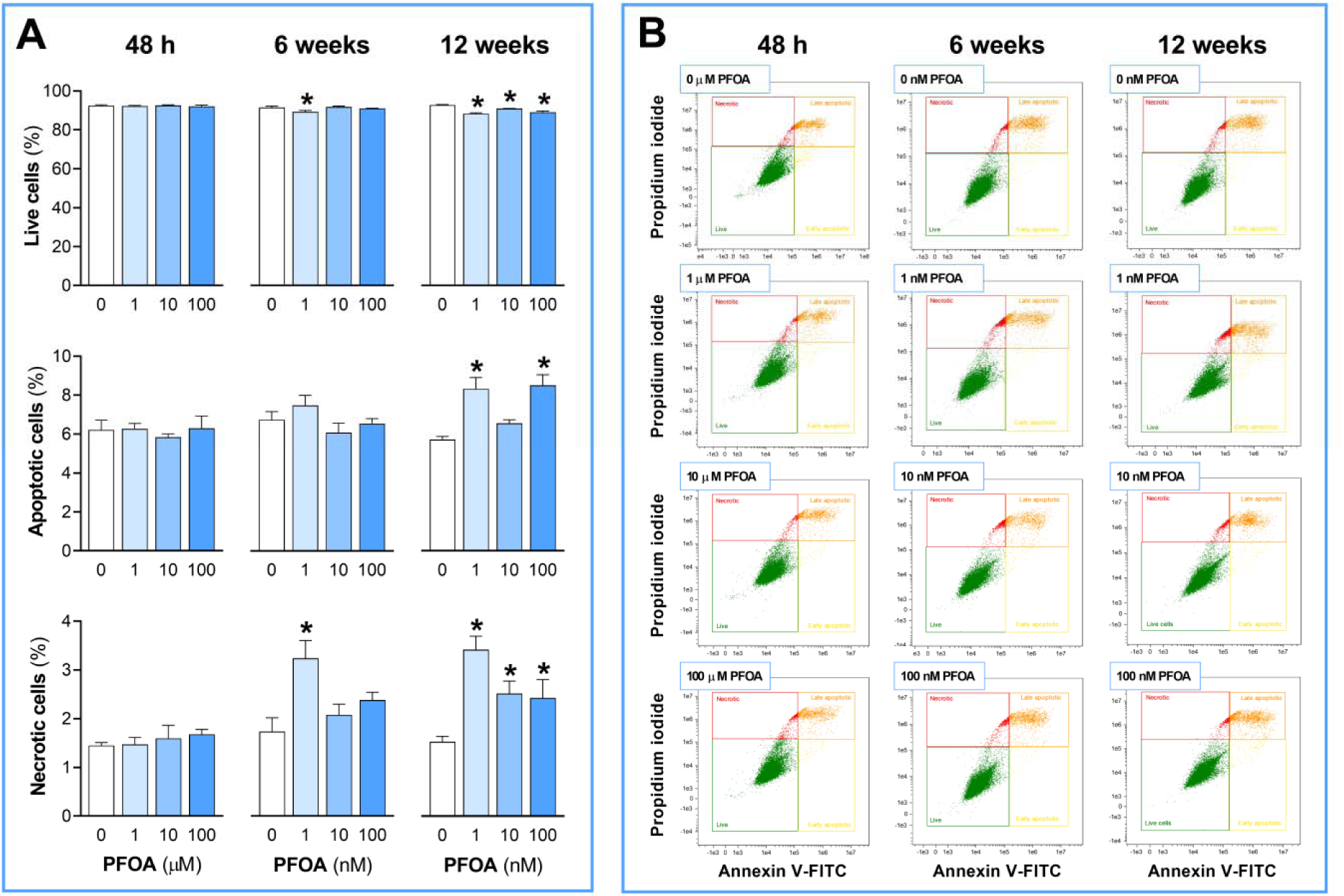
Effect of short-term and long-term exposure of EA.hy926 cells to PFOA on apoptosis and necrosis. EA.hy926 cells were exposed to 1 μM, 10 μM, and 100 μM PFOA for 48 h (short-term exposure) and 1 nM, 10 nM, and 100 nM PFOA for 6 and 12 weeks (long-term exposure), followed by the assessment of apoptosis and necrosis using Annexin V-FITC and propidium iodide on a flow cytometer. **(A)** Results are expressed as the percentage of live, apoptotic, and necrotic cells. Each data bar represents the mean ± SEM of 4 independent experiments (short-term exposure) or 3 cell culture flasks (long-term exposure). **(B)** Representative dot-plots from flow cytometry are shown. \**p* <0.05 *vs.* control.

#### 3.1.2 Cell cycle

Exposure of EA.hy926 cells to PFOA for 48 h and 6 weeks had no impact on cell cycle progression across any of the tested concentrations. However, after 12 weeks of exposure to 100 nM PFOA, we observed a decrease in the percentage of cells in the G_0_/G_1_ phase and a corresponding increase in the S phase, but the percentage of cells in the G_2_/M phase remained comparable to the control group. Notably, the percentage of cells in the G_2_/M phase was significantly elevated following 12 weeks of exposure to 10 nM PFOA (Figure 2A and B).

**Figure 2.**
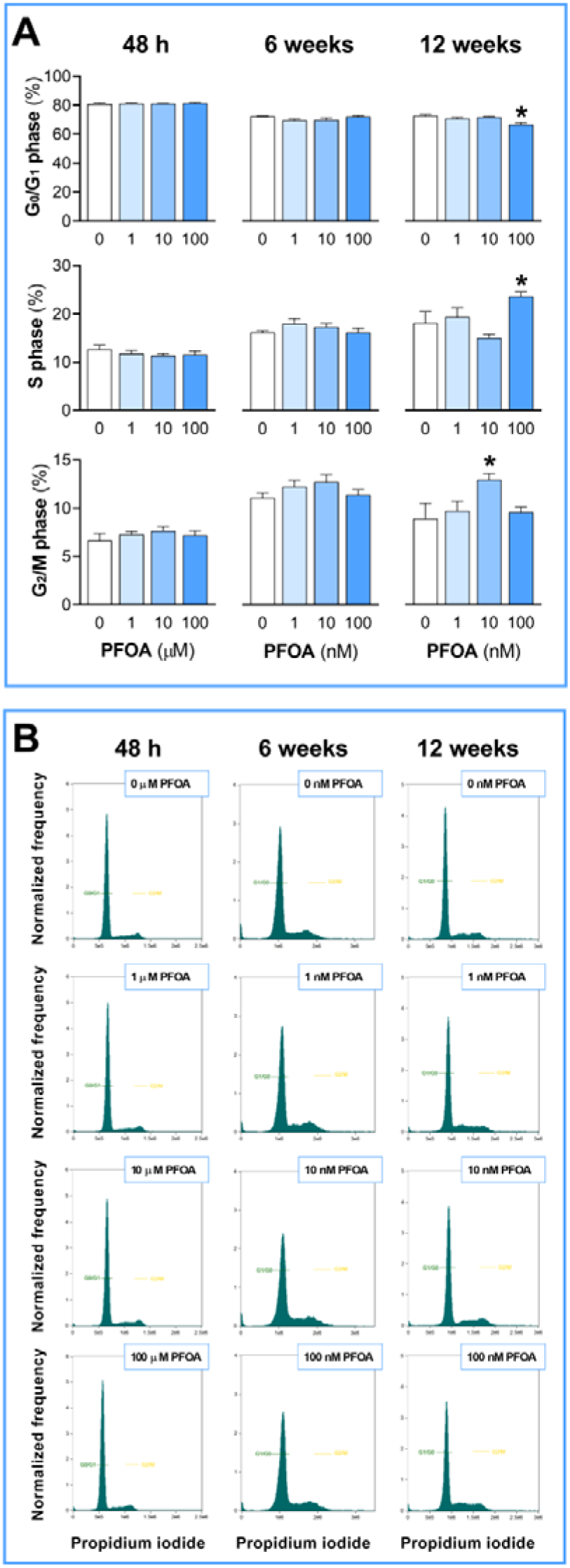
Effect of short-term and long-term exposure of EA.hy926 cells to PFOA on cell cycle. EA.hy926 cells were exposed to 1 μM, 10 μM, and 100 μM PFOA for 48 h (short-term exposure) and 1 nM, 10 nM, and 100 nM PFOA for 6 and 12 weeks (long-term exposure), followed by cell cycle analysis using propidium iodide on a flow cytometer. **(A)** Results are expressed as the percentage of cells in G_0_/G_1_, S, and G_2_/M phase. Each data bar represents the mean ± SEM of 4 independent experiments (short-term exposure) or 3 cell culture flasks (long-term exposure). **(B)** Representative histograms from flow cytometry are shown. \**p* <0.05 *vs.* control.

#### 3.1.3 Endothelial permeability and monocyte adhesion to endothelial cell monolayer

A significant increase in endothelial permeability was noted only after 12 weeks of exposure of EA.hy926 cells to 1 nM, 10 nM, and 100 nM of PFOA (Figure 3A), whereas monocyte adhesion to the endothelial cell monolayer significantly increased after 6 weeks of exposure to 10 nM PFOA and after 12 weeks of exposure to 100 nM PFOA (Figure 3B and C).

**Figure 3.**
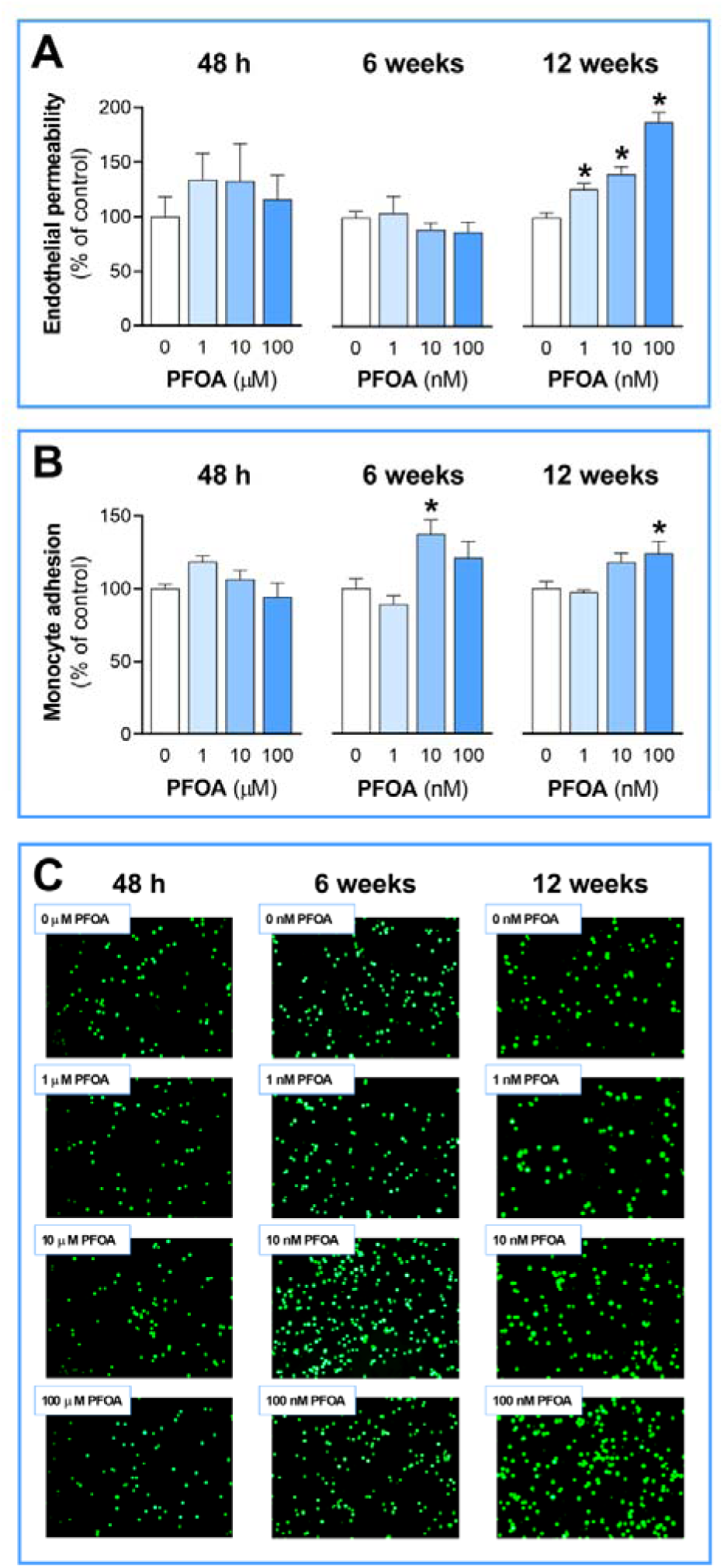
Effect of short-term and long-term exposure of EA.hy926 cells to PFOA on endothelial permeability and monocyte adhesion to endothelial cell monolayer. EA.hy926 cells were exposed to 1 μM, 10 μM, and 100 μM PFOA for 48 h (short-term exposure) and 1 nM, 10 nM, and 100 nM PFOA for 6 and 12 weeks (long-term exposure), followed by the assessment of **(A)** endothelial permeability using the two-compartment permeability assay with FITC-labeled dextran and **(B)** the extent of monocyte adhesion to the EA.hy926 monolayer using calcein AM-labeled U937 cells. **(A, B)** Results were expressed relative to the vehicle-treated control (100%). Each data bar represents the mean ± SEM of 4 independent exper iments (short-term exposure) or 3 cell culture flasks (long-term exposure). **(C)** Representative fluorescent photomicrographs of calcein AM-labeled U937 cells adhered to the EA.hy926 cell monolayer are shown. \**p* <0.05 *vs.* control.

#### 3.1.4 Cell migration and endothelial tube formation

We next examined angiogenesis-related processes, including endothelial cell adhesion to the extracellular matrix, cell migration, and endothelial tube formation. Both short-term and long-term exposure of EA.hy926 cells to PFOA did not affect adhesion to fibronectin at any of the tested concentrations (Supplementary Figure 3). However, cell migration increased after 6 weeks of exposure to 10 nM and 100 nM PFOA but decreased after 12 weeks of exposure to 100 nM PFOA (Figure 4A, Supplementary Figure 4A). Endothelial tube formation was affected only after 12 weeks of exposure to 10 nM and 100 nM PFOA. We observed a significant increase in the total length of tubes, as well as a higher number of nodes and junctions, and greater total branching length, indicating enhanced endothelial tube formation (Figure 4B, Supplementary Figure 4B). Additionally, we investigated ROS production following both short-term and long-term exposure to PFOA. The results showed significantly reduced ROS levels after 6 weeks of exposure to 10 nM and 100 nM PFOA (Supplementary Figure 5).

**Figure 4.**
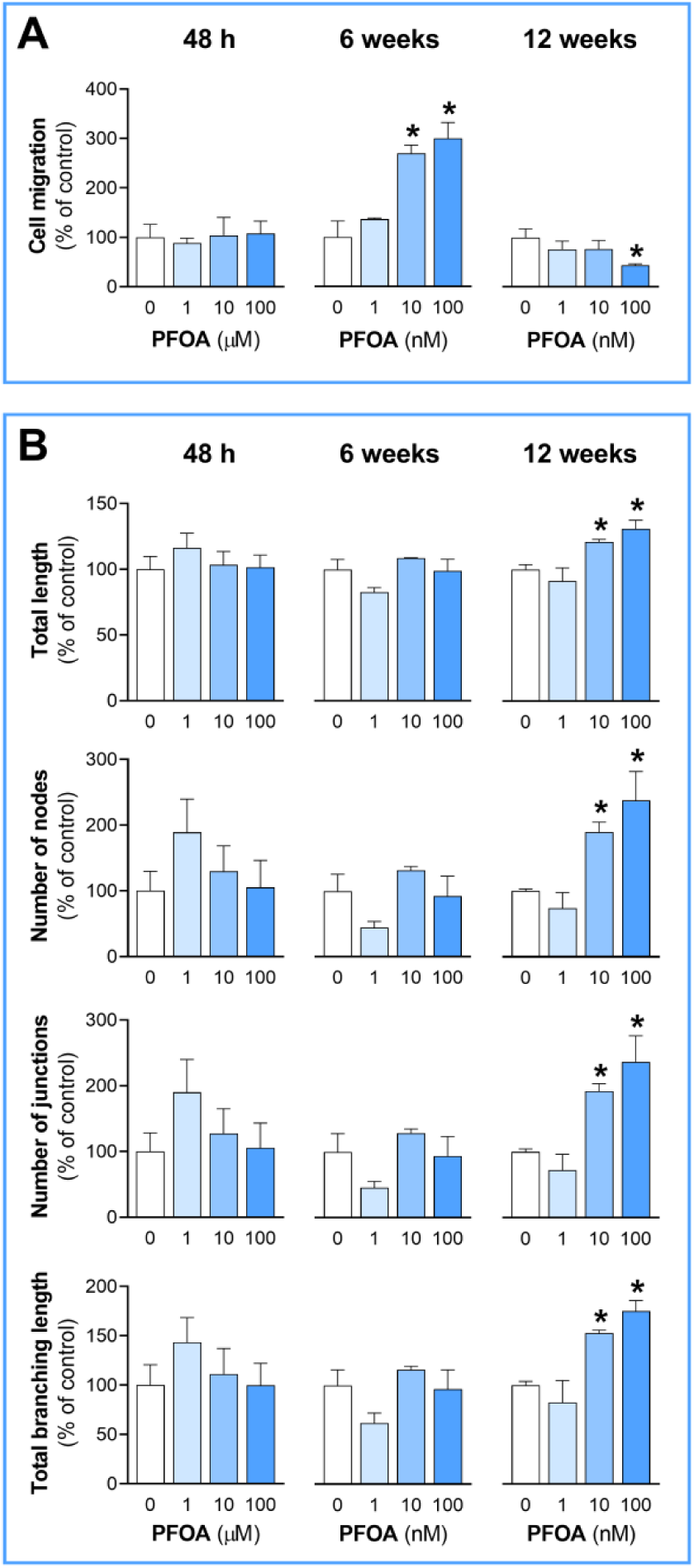
Effect of short-term and long-term exposure of EA.hy926 cells to PFOA on cell migration and endothelial tube formation. EA.hy926 cells were exposed to 1 μM, 10 μM, and 100 μM PFOA for 48 h (short-term exposure) and 1 nM, 10 nM, and 100 nM PFOA for 6 and 12 weeks (long-term exposure), followed by the assessment of **(A)** cell migration using the modified Boyden chamber assay and **(B)** the extent of angiogenesis *in vitro* using three-dimensional matrix Cultrex BME. Results were expressed relative to the vehicle-treated control (100%). Each data bar represents the mean ± SEM of 4 independent experiments (short-term exposure) or 3 cell culture flasks (long-term exposure). \**p* <0.05 *vs.* control.

### 3.2 Identification of differentially expressed genes

Whole-genome transcriptome analysis identified differentially expressed genes (DEGs) in EA.hy926 cells, with at least a 2-fold change and a Q value of ≤0.05, following both short-term and long-term exposure to specified concentrations of PFOA (Figure 5A). The number of DEGs varied depending on the duration of exposure and the concentration of PFOA. After 48 h, the highest number of DEGs was observed at the highest concentration of 100 µM PFOA, resulting in 359 DEGs (119 upregulated, 240 downregulated). In contrast, exposure to 10 µM PFOA and 1 µM PFOA resulted in 197 DEGs (88 upregulated, 109 downregulated) and 98 DEGs (61 upregulated, 37 downregulated), respectively. Interestingly, the greatest overall number of DEGs was detected after 6 weeks of exposure to 1, 10, and 100 nM PFOA, surpassing the counts from 48-h exposure to the higher micromolar concentrations and the 12-week exposure to the same nanomolar concentrations of PFOA. Notably, exposure to 10 nM PFOA for 6 weeks produced the highest total of 6,018 DEGs, comprising 5,038 upregulated and 980 downregulated genes. Conversely, exposure to 1 nM and 100 nM PFOA yielded 397 DEGs (311 upregulated, 86 downregulated) and 3,811 DEGs (3,504 upregulated, 307 downregulated), respectively. A similar trend was evident after 12 weeks of exposure to the three nanomolar concentrations of PFOA. Again, the highest number of DEGs was associated with 10 nM PFOA, with a total of 1,725 DEGs (1,627 upregulated, 98 downregulated). In comparison, exposure to 1 nM PFOA and 100 nM PFOA resulted in 205 DEGs (109 upregulated, 96 downregulated) and 904 DEGs (701 upregulated, 203 downregulated), respectively (Figure 5B).

**Figure 5.**
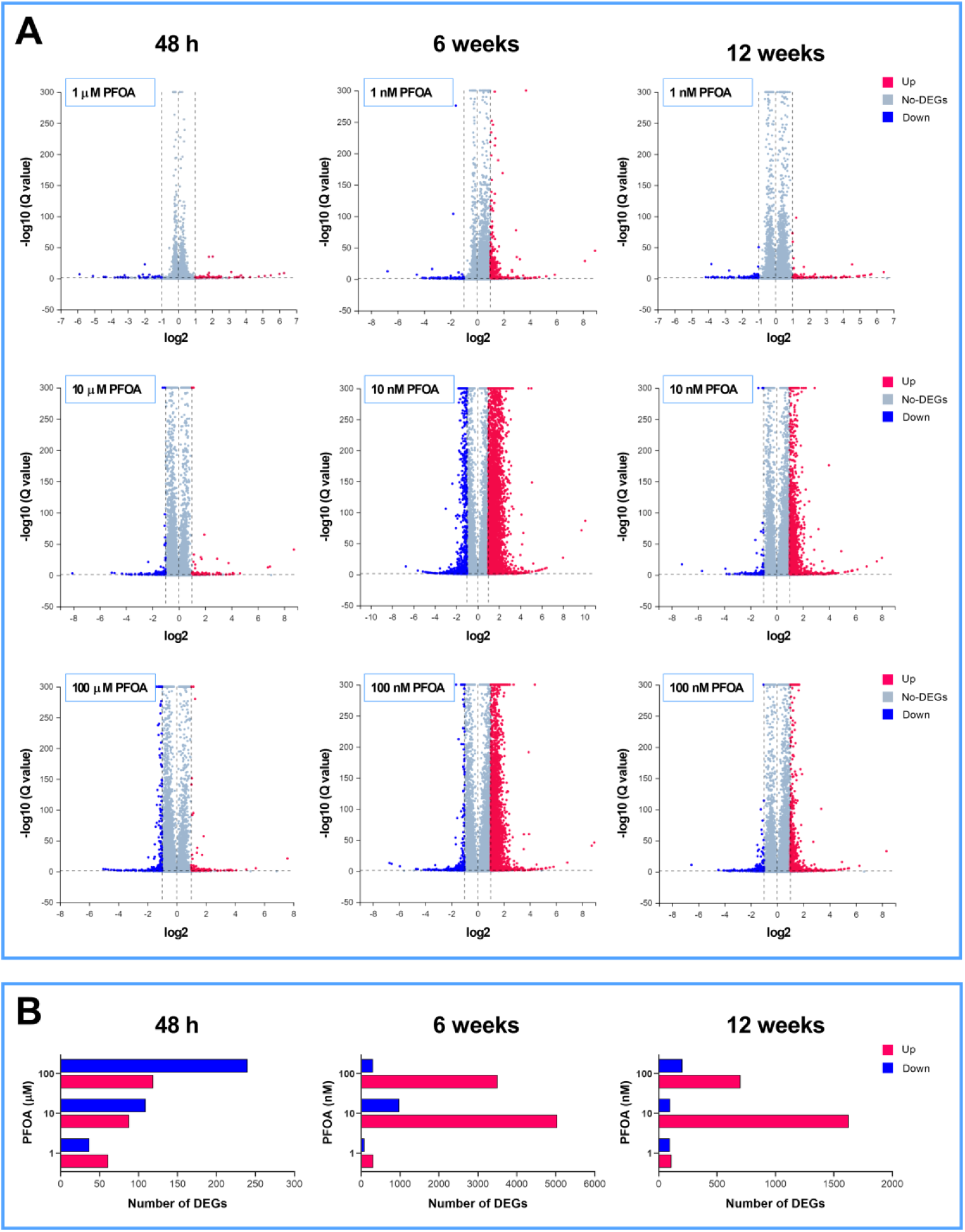
Volcano scatter plots depicting global mRNA expression and the number of differentially expressed genes (DEGs) following short-term (48 h) and long-term (6 and 12 weeks) exposure of EA.hy926 cells to specified concentrations of PFOA. **(A)** The X-axis represents the fold change of the difference after conversion to log2, whereas the Y-axis represents the significance value (Q value) after conversion to –log10. In each plot, significantly upregulated genes are marked as red dots, downregulated genes as blue dots, and non-significant findings as gray dots. **(B)** The X-axis displays the number of DEGs, and the Y-axis shows the different concentrations of PFOA. For each concentration, red bars represent the number of upregulated genes, while blue bars represent the number of downregulated genes.

The results presented in Supplementary Figure 6 illustrate the hierarchical clustering of DEGs, highlighting distinct expression patterns associated with short-term (48 h) and long-term (6 and 12 weeks) exposure to micromolar and nanomolar concentrations of PFOA, respectively. Figure 6 details the number of unique and overlapping DEGs following short-term and long-term exposure of EA.hy926 cells to selected PFOA concentrations. The Venn diagram demonstrates that exposure to different micromolar concentrations of PFOA resulted in only 10 shared DEGs after 48 h. In contrast, the number of shared genes among the three nanomolar concentrations of PFOA peaked after 6 weeks, reaching a total of 293 DEGs. After 12 weeks of exposure, 55 shared DEGs were identified.

**Figure 6.**
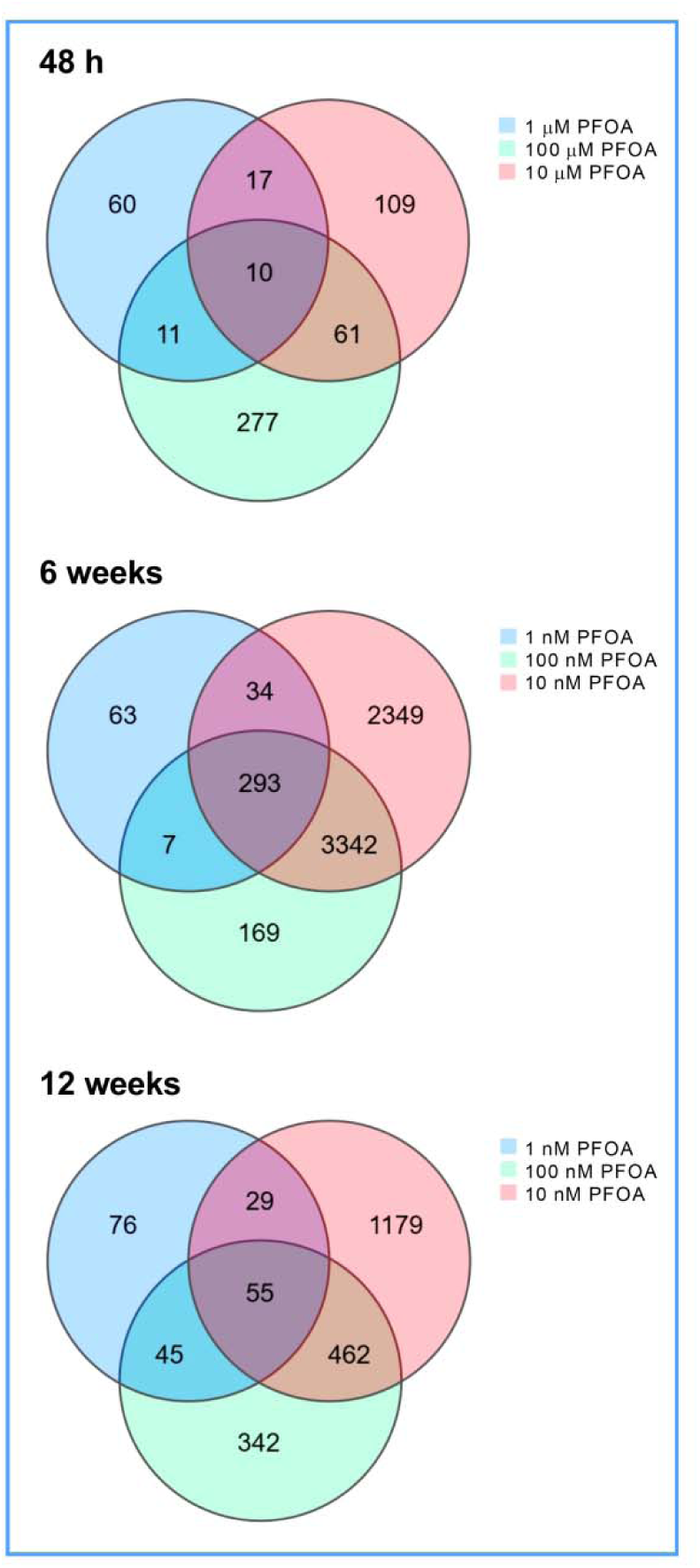
Venn diagram illustrating differentially expressed genes (DEGs) after short-term (48 h) and long-term (6 and 12 weeks) exposure of EA.hy926 cells to PFOA. The diagram highlights the number of unique and overlapping DEGs among the specified concentrations of PFOA.

### 3.3 Determination of benchmark concentration

To visualize changes in response to increasing concentrations of PFOA over different exposure durations (short-term – 48 h and long-term – 6 and 12 weeks) and levels of biological organization (apical – A and transcriptomics – T), we plotted accumulation curves for the BMC values (Figure 7). The accumulation plots across three time points and two biological levels displayed similar curve shapes, with the initial segments exhibiting shallow slopes across the BMCs, until reaching a phase referred to as “initiation” or “concentrated molecular response” (Johnson et al., 2022). Notably, after 48 h, A-POD did not show a concentration-dependent response to PFOA exposure.

**Figure 7.**
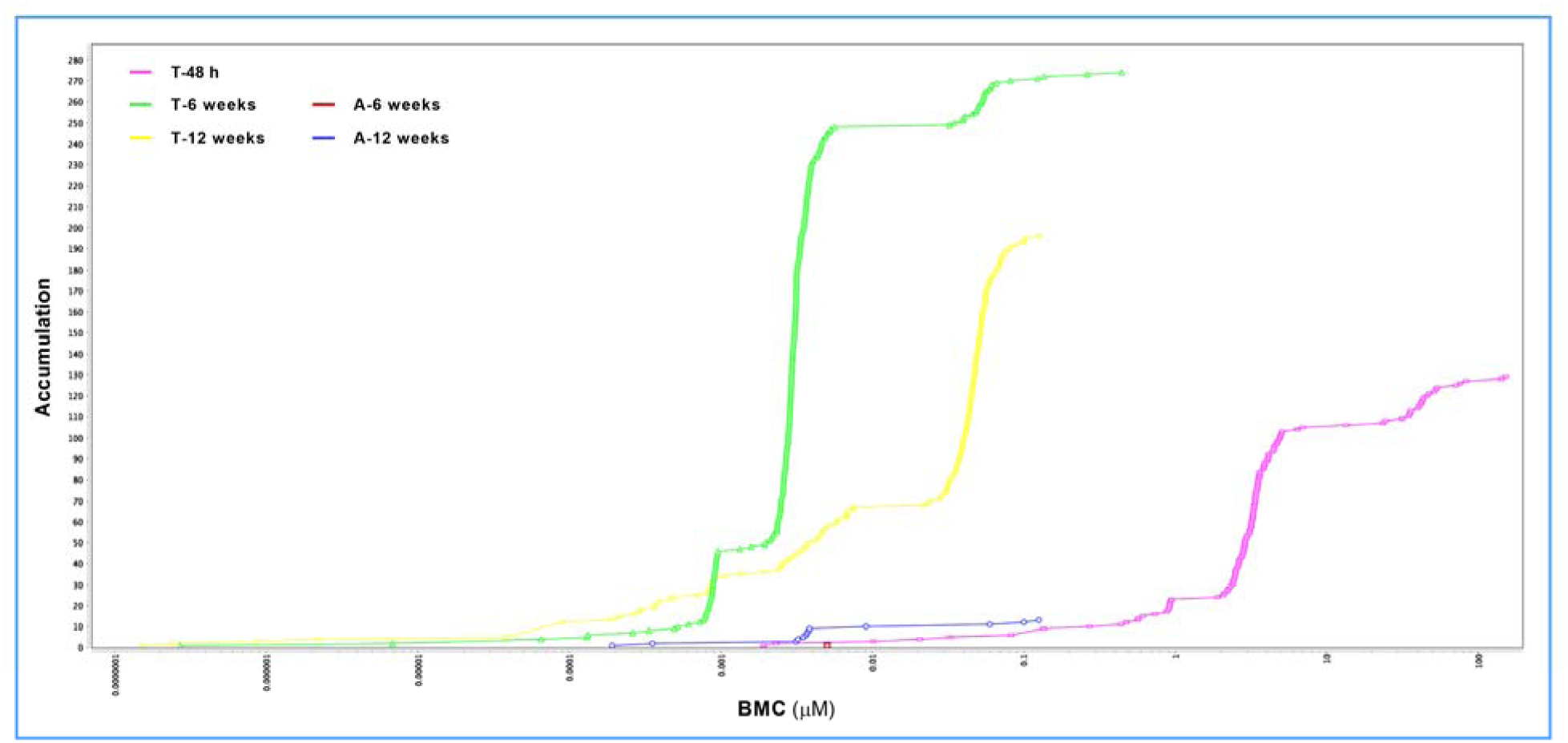
Accumulation plots of benchmark concentration (BMC) values for apical (A) and transcriptomics (T) data following short-term (48 h) and long-term (6 and 12 weeks) exposure of EA.hy926 cells to PFOA. The analysis was conducted in BMDExpress v3.20.0106.

The calculated average BMC values were as follows: 5 nM for A-6 weeks, 8.6 nM for A-12 weeks, 10.3 µM for T-48 h, 6.8 nM for T-6 weeks, and 36.8 nM for T-12 weeks. The average BMCL and BMCU values were determined as follows: for A-6 weeks, 3.7 nM and 12.1 nM; for A-12 weeks, 5.4 nM and 28.2 nM; for T-48 h, 6.3 µM and 36.8 µM; for T-6 weeks, 4.1 nM and 19.9 nM; and for T-12 weeks, 22.1 nM and 134 nM; respectively. Notably, the BMCL value (POD) for T-6 weeks was 5.5 times lower than that for T-12 weeks and ∼1,500 times lower than that for T-48 h. The POD values for A-6 weeks and A-12 weeks were comparable to that of T-6 weeks but were 6 times and 4.1 times lower than the POD value for T-12 weeks, respectively (Figure 8).

**Figure 8.**
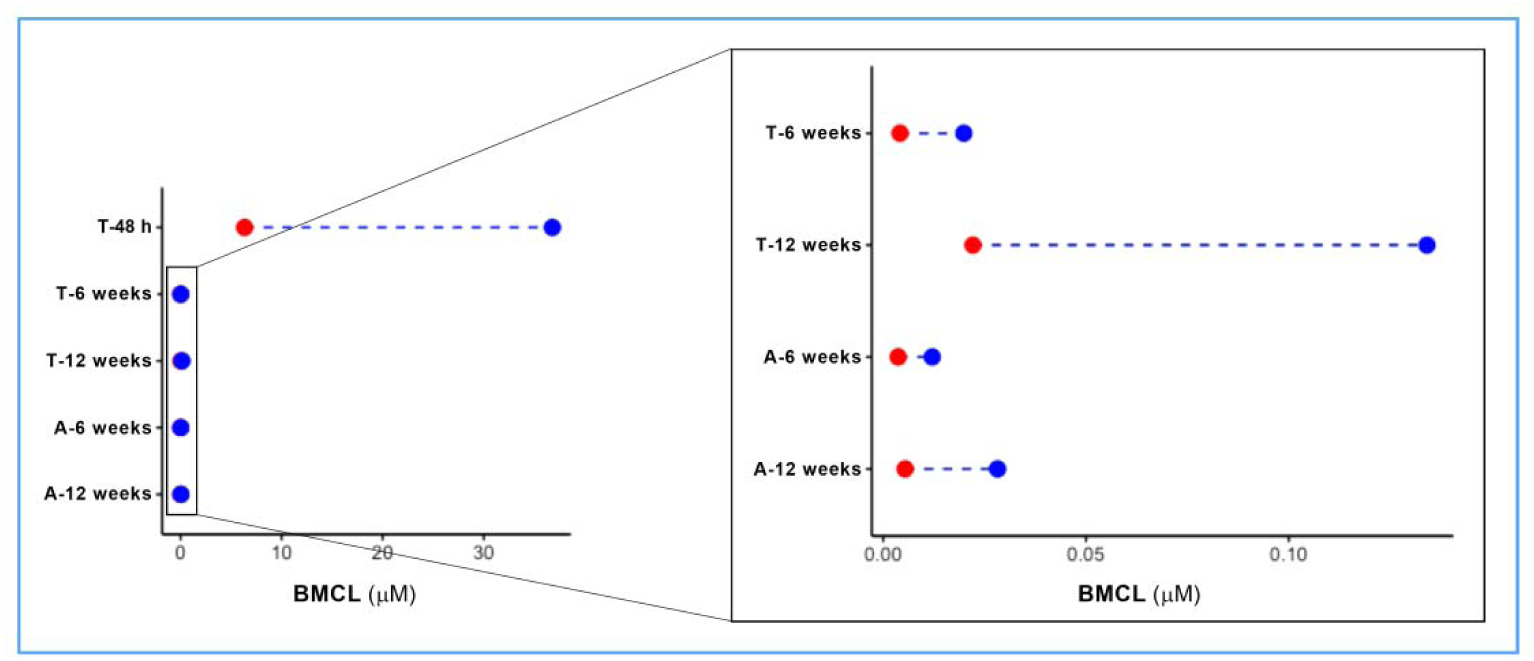
Range plots of benchmark concentration lower confidence limit (BMCL; red dot) and benchmark concentration upper confidence limit (BMCU; blue dot) values for apical (A) and transcriptomics (T) data following short-term (48 h) and long-term (6 and 12 weeks) exposure of EA.hy926 cells to PFOA. For improved visualization, the right panel shows BMCL and BMCU values for all time points except T-48 h.

The density plot curves of BMC indicate unimodal distributions for T-48 h and T-6 weeks, suggesting that the BMC values were positively skewed. In contrast, the T-12 weeks curve was bimodal, revealing two distinct subgroups of BMC values (Figure 9).

**Figure 9.**
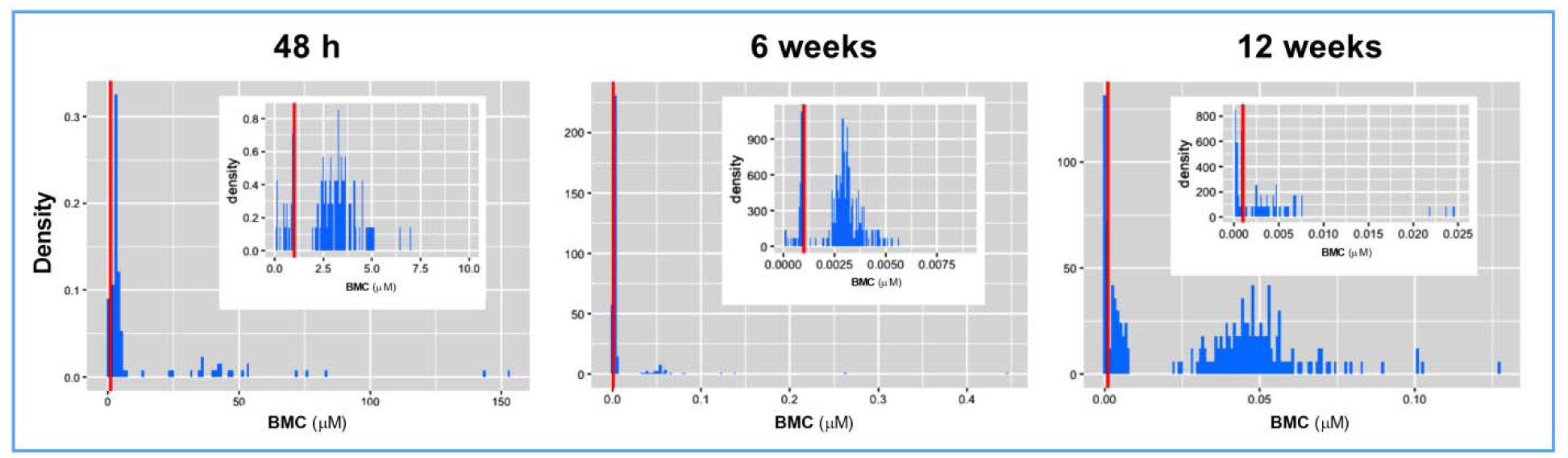
Density plots illustrating the distribution of benchmark concentration (BMC) values following short-term (48 h) and long-term (6 and 12 weeks) exposure of EA.hy926 cells to PFOA. For improved visualization, the smaller panel within the main panel shows a specific narrow range of BMC values.

DAVID functional analysis was conducted using transcriptomics BMCL values to evaluate the functional modules affected by both short-term and long-term exposure of EA.hy926 cells to PFOA. The results of the functional enrichment analysis revealed that genes in the T-48 h and T-6 weeks groups share several functional modules related to transcription, including “Regulation of transcription by RNA polymerase II,” “DNA-binding,” and “DNA-binding transcription factor activity, RNA polymerase II specific”. The functional module with the lowest estimated BMCL value in these two groups was the interleukin-17 (IL-17) signaling pathway, with average T-POD values close to the 25^th^ percentile. In contrast, the functional modules associated with the genes affected by 12-week-long exposure to PFOA (T-12 weeks) differed from those identified in the T-48 h and T-6 weeks groups. These included “Extracellular space,” “Extracellular matrix,” “Cytokine-cytokine interaction,” and “Cell adhesion”. Notably, the functional modules with the lowest estimated BMCL related to endothelial dysfunction in the T-12 weeks group were “Cell adhesion” and “Positive regulation of gene expression” (Figure 10).

**Figure 10.**
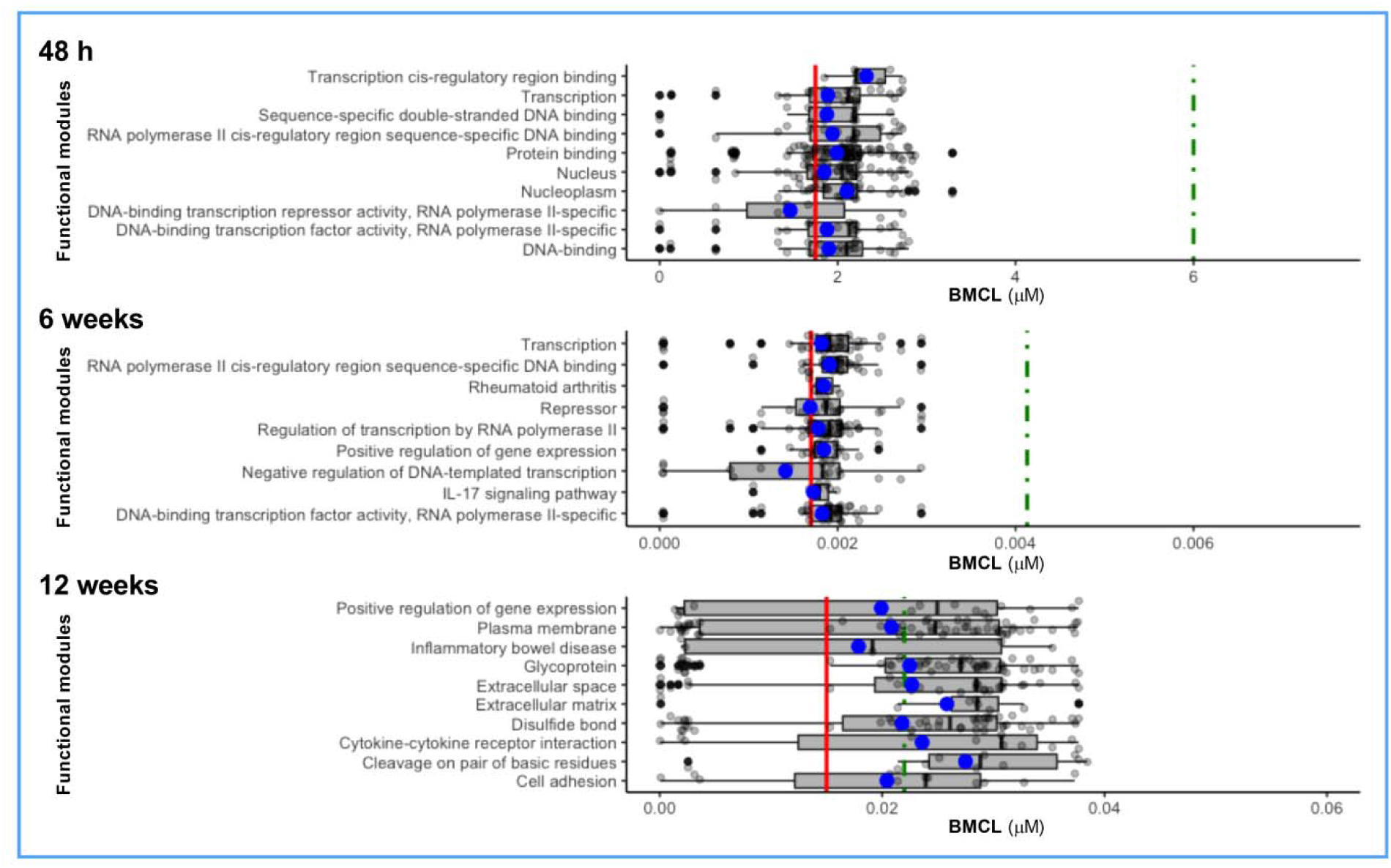
Benchmark concentration lower confidence limit (BMCL) values of individual genes classified into functional modules. The analysis of functional modules was performed using the DAVID bioinformatics tool. The grey box represents the range between the 1^st^ and the 3^rd^ quartiles, while the thick black line indicates the mean. The red vertical line marks the lowest 25^th^ percentile of BMCL, and the green dotted vertical line indicates the median BMCL value at each exposure time point.

## 4. Discussion

The use of *in vivo* apical assays to assess dose-response relationships following chemical exposure has been a hallmark of chemical risk assessment for decades. However, this approach has notable limitations, prompting the exploration of replacement methods known as new approach methodologies (NAMs). NAMs, including toxicogenomics, have emerged as a promising tool for future alternative testing paradigms that could address some of the drawbacks associated with traditional apical testing. An alternative to *in vivo* traditional chemical risk assessment has been proposed by the United States National Research Council, the Tox21 Program, and the European Commission’s Advisory Group for Endocrine Disruptors. This approach emphasizes the importance of using human cells *in vitro*, which can enhance the relevance of results to human health while addressing several limitations of *in vivo* studies (Krewski et al., 2010). When combined with BMC modeling, *in vitro* methods utilizing human cells may provide a viable alternative to *in vivo* experiments, thereby advancing research in human health risk assessment.

Several studies have employed *in vitro* exposure of human cells to derive POD values for a variety of chemicals (Addicks et al., 2023; Matteo et al., 2023; Reardon et al., 2021), providing strong support for the use of *in vitro* assays in evaluating the potency of various chemicals to impact human health. However, important issues remain unresolved, including the selection and use of different levels of biological organization and exposure time points in chemical hazard characterization. Comparisons across different biological levels and exposure durations have been made in several animal studies published to date. It has been shown that transcriptomics-based BMD and the associated T-POD values from liver and kidney typically fall within a factor of 5 of the lowest apical (histopathological) BMD value, suggesting that the transcriptomics-based BMD could be the method of choice for risk assessment *in vivo* (Gwinn et al., 2020). Additionally, it has been reported that *in vivo* short-term toxicogenomics testing was sufficiently sensitive to serve as an alternative to chronic *in vivo* studies for deriving PODs for triazole fungicides (LaRocca et al., 2020). A similar concordance was observed in fathead minnow larvae, where liver, kidney, and whole animal transcriptomics data from short-term exposure studies aligned with apical effects from long-term studies (Martin et al., 2023). The concordance between sub-chronic T-POD and chronic A-POD supports the use of sub-chronic transcriptomics-derived PODs for estimating A-POD values within a risk-based framework for chronic toxicity and carcinogenicity (Bianchi et al., 2021). The study by Bhat and colleagues demonstrated the superiority of the transcriptomics approach over the apical approach in evaluating conazoles. The authors found that T-PODs following a 30-day exposure accurately ranked four tumorigenic and one non-tumorigenic conazole by their tumorigenic potency, which was not possible with the corresponding 30-day apical assay (liver weight) BMD and A-POD data. Furthermore, the 30-day T-PODs for the four tumorigenic conazoles were generally more sensitive than the 2-year A-POD values for hepatocellular carcinoma and adenoma (Bhat et al., 2013). It appears that in rodents, short-term T-POD values are at least as sensitive as long-term A-POD values, making them a suitable alternative to traditional chemical risk assessment. Similar findings were observed in fish exposed to estrogenic chemicals, where short-term T-PODs could directly substitute for chronic toxicity PODs (Pagé-Larivière et al., 2019). However, all the above-mentioned data were obtained from animal studies and may not accurately reflect the sensitivity of various biological levels or exposure durations in human cells. The results of this study provide a certain degree of deflection from the traditional animal studies by presenting novel information regarding the sensitivity of transcriptomics and apical approaches in different exposure contexts involving human cells. The main finding of the current study is that *in vitro* chronic T-POD and A-POD values are more sensitive than acute T-POD values in human endothelial cells. Notably, acute A-POD could not be determined, as none of the apical measurements showed a concentration-dependent response upon exposure to micromolar concentrations of PFOA. The difference in the range of PODs between the short-term and long-term exposures suggests that acute T-POD may not be the best estimate for chronic A-POD in PFOA-exposed human endothelial cells. This implies a shift in POD values between short-term and long-term exposure, indicating that the potency of PFOA to alter the function of human endothelial cells depends on the duration of exposure. Similar findings were reported in the study of Addicks and colleagues on human liver spheroids, where the potency of nearly all investigated individual PFAS and their mixtures increased from 24 h to 10 days, with the exception of one mixture and PFOS (Addicks et al., 2023). In our study, the concentrations of PFOA used for long-term exposure were 1,000 times lower than those used for short-term exposure. However, the BMD approach does not depend on dose selection, as demonstrated in the determination of NOAEL (Davis et al., 2011). We adjusted the range of PFOA concentrations for long-term experiments because high concentrations of chemicals in prolonged exposure scenarios can elicit more robust toxicogenomic responses, potentially leading to an underestimation of T-POD (Pagé-Larivière et al., 2019). These findings highlight the importance of considering both concentration and exposure duration when designing *in vitro* toxicogenomic experiments to determine PODs (Carpi et al., 2024).

We also analyzed the BMCs across different biological levels, contributing to the notion that molecular analysis can identify initial events leading to effects at higher levels of organization (Martinez et al., 2020). Although studies investigating this concept are limited, those that exist indicate that similar slopes in cumulative effect plots from the BMD analysis – derived from transcriptomics, metabolomics, and apical (morphometric) data following exposure to tributyltin (Martinez et al., 2020) and PFOS (Martinez et al., 2019) – reflect the propagation of molecular events from the transcriptome and metabolome to higher organizational levels such as morphology. In this study, comparisons of BMDs across different biological levels (transcriptome and apical) provided some, but not definitive, mechanistic connections. The difference between the acute T-POD and chronic A-POD (cell migration after 6 weeks, and angiogenesis and endothelial permeability after 12 weeks of exposure of EA.hy926 cells to PFOA) suggests that transcriptomics data obtained after short-term exposure of EA.hy926 cells to PFOA may not serve as an early warning sign for effects at higher levels of biological organization that arise from prolonged exposure. Functional module analysis using genes with calculated PODs after 48 h revealed that gene transcription could be a key molecular event through which PFOA affects the function of human endothelial cells. While this molecular pathway may relate to observed apical changes after 6 and 12 weeks of exposure to PFOA, it could also intersect with various biological processes in human endothelial cells. Although we considered the differences between short-term T-POD and long-term A-POD as a primary limitation in linking early molecular changes to later alterations at higher biological levels, the lack of an early warning sign may be specific to human endothelial cells, which is a notion that should be further explored using other *in vitro* human cell models.

The connections between different biological levels were more pronounced in long-term exposure scenarios. After 6 weeks of exposure, we observed enrichment of specific molecular events closely linked to the apical data, particularly the IL-17 signaling pathway, which could be the molecular event responsible for cell migration. It has been shown that IL-17 promotes the migration and invasion of human cancer cells (Guo et al., 2019) and trophoblasts (Zhang et al., 2022). After 12 weeks of exposure, enrichment analysis of genes with calculated PODs revealed additional specific molecular events associated with endothelial function, including extracellular matrix interactions and cytokine-cytokine interactions. There is substantial evidence linking these processes to angiogenesis (Davis and Kemp, 2023; Geindreau et al., 2022; Mongiat et al., 2016; Neve et al., 2014; Wang et al., 2023) and endothelial permeability (Alexander and Elrod, 2002; Hendel et al., 2014; Sluiter et al., 2021). Therefore, similar BMC values in *in vitro* long-term exposure studies may reflect the propagation of molecular events (transcriptome) to higher levels of organization (apical).

## 5. Conclusions

Selecting appropriate time points and biological outputs poses a significant challenge in toxicological studies. Our primary objective in this study was to investigate the sensitivity of *in vitro* transcriptomics and apical assays across various exposure scenarios using BMC modeling. Our findings indicate that, in PFOA-exposed human endothelial cells, chronic T-POD is a more superior approach for hazard characterization than acute T-POD. The most sensitive molecular pathway affected by acute, short-term exposure to PFOA primarily relates to general toxicity. In contrast, pathways affected by chronic, long-term exposure serve as early molecular indicators of toxicity at higher levels of biological organization, highlighting the importance of mechanistically identifying initial toxic events. This work contributes to the growing understanding of the human health hazards associated with PFOA exposure and underscores the use of chronic *in vitro* exposure scenarios and BMC modeling as a more sensitive approach for chemical risk assessment. While this study was conducted using human endothelial cells to provide more relevant data, it is important to note that these findings may be specific to this cell type. Investigating other human-derived, toxicologically relevant cells would further enhance our understanding of the selection and applicability of different approaches for chemical risk assessment.

## Data availability

Data will be made available on request.

