## Supplementary Material for "Comparative sensitivity of *in vitro* acute and chronic apical and transcriptomic points of departure for perfluorooctanoic acid in human vascular endothelial cells"

**SUPPLEMENTARY MATERIAL AND METHODS**

**Cell viability assay**

The alamarBlue^TM^ assay was utilized to quantify relative cytotoxicity following short-term and long-term exposure of EA.hy926 cells to PFOA. For short exposure, EA.hy926 cells were plated in 96-well plates (2.5 × 10^4^ cells per well; 6 wells per each treatment) and exposed the following day to either vehicle (0.05% DMSO) or 1 μM, 10 μM, and 100 μM PFOA for 48 h. In long-term exposure experiments, EA.hy926 cells were harvested from each flask after 6 and 12 weeks of continuous exposure to either vehicle (0.05% DMSO) or 1 nM, 10 nM, and 100 nM PFOA, plated in 96-well plates (2.5 × 10^4^ cells per well; 6 wells per each flask), and exposed to the corresponding treatments for an additional 48 h. The alamarBlue^TM^ assay was performed following the protocol published elsewhere (Kokai et al., 2020). Quantification of the obtained results involved calculating the mean fluorescent signal for each treatment and control. The metabolic activity of PFOA-exposed EA.hy926 cells was then expressed as a percentage of the metabolic activity relative to the control, which was set at 100%.

**Apoptosis assay**

The apoptosis assay using Annexin V-FITC and propidium iodide (PI) was performed to quantitatively determine the percentage of EA.hy926 cells undergoing apoptosis and necrosis following both short-term and long-term exposure to PFOA. For short-term exposure, EA.hy926 cells were plated in 35-mm dishes (0.7 × 10^6^ cells per dish; two dishes per treatment) and exposed the following day to either vehicle (0.05% DMSO) or 1 μM, 10 μM, and 100 µM PFOA for 48 h. After exposure, the growth medium was collected, and cells were lifted into suspension using a 0.25% trypsin-EDTA solution for 4 min. Complete culture medium was then added, and the cells were centrifuged at 400 × g for 5 min. The supernatant was carefully removed, and the cell pellet was resuspended in fresh complete culture medium. The cells were counted, and 1.5 × 10^6^ cells (both live and dead) were transferred to 2 mL microcentrifuge tubes for a second centrifugation at 400 × g for 5 min. In long-term exposure experiments, EA.hy926 cells (a total of 1.5 × 10^6^ cells per flask, both live and dead) and their growth medium were harvested from each flask after 6 and 12 weeks of continuous exposure to either vehicle (0.05% DMSO) or 1 nM, 10 nM, and 100 nM PFOA into 15 mL tubes by trypsinization, followed by centrifugation at 400 × g for 5 min. In both short-term and log-term experiments, the supernatant was discarded, and the cell pellet was washed with 2 mL of PBS per tube. The cell suspension in PBS was centrifuged again at 400 × g for 5 min, after which the pellet was resuspended in 50 µL of Annexin V Binding Buffer (10 mM HEPES, 140 mM NaCl, 2.5 mM CaCl_2_, pH 7.4) containing 2.5 µL of commercially obtained Annexin V-FITC Reagent and 2.5 µL of PI solution (100 µg/mL). Following a 20-min incubation in the dark, 200 µL of Annexin V Binding Buffer was added to each tube, and the mixture was filtered through a nylon mesh into 1.5 mL tubes. The tubes were kept on ice and protected from light until analysis. Cells were analyzed using the Amnis® ImageStream®X Mk II Imaging Flow Cytometry (IFC) (Luminex Corporation, Austin, USA) and the INSPIRE™ ImageStream®X System Software (Luminex Corporation, Austin, USA). Parameters were set to detect individual cells, targeting an optimal count of 25,000 cells, using a blue laser (excitation wavelength at 488 nm) with 100 mW power for the FITC and PI channels. The combination of flow cytometry and microscopy in the Amnis® ImageStream®X Mk II IFC enables visual verification of results, precise placement of regions of interest, and reliable conclusions. The data analysis was conducted using the IDEAS® 6.2 Image Data Exploration and Analysis Software (Luminex Corporation, Austin, USA), employing dot-plot representation and four defined quadrants based on fluorescent signal intensity from FITC and PI. Results were presented as the percentage of cells in each quadrant, specifically indicating the proportions of live, apoptotic (both early and late), and necrotic cells.

**Cell cycle analysis**

Flow cytometry using PI was employed to quantitatively assess the percentage of EA.hy926 cells in different phases of the cell cycle following both short-term and long-term exposure to PFOA. For short-term exposure, EA.hy926 cells were plated in 35-mm dishes (0.7 × 10^6^ cells per dish; two dishes per treatment) and exposed the following day to either vehicle (0.05% DMSO) or 1 μM, 10 μM, and 100 µM PFOA for 48 h. After exposure, cells were lifted into suspension using a 0.25% trypsin-EDTA solution for 4 min. Complete culture medium was then added, and the cells were centrifuged at 400 × g for 5 min. The supernatant was carefully removed, and the cell pellet was resuspended in fresh complete culture medium. The cells were counted, and 1.5 × 10^6^ cells were transferred to 15 mL tubes for a second centrifugation at 400 × g for 5 min. In long-term exposure experiments, EA.hy926 cells were harvested from each flask (1.5 × 10^6^ cells per flask) after 6 and 12 weeks of continuous exposure to either vehicle (0.05% DMSO) or 1 nM, 10 nM, and 100 nM PFOA into 15 mL tubes by trypsinization, followed by centrifugation at 400 × g for 5 min. In both short-term and log-term experiments, the supernatant was discarded, and the cell pellet was washed with 5 mL of ice-cold PBS per tube. The cell suspension in PBS was then centrifuged again at 400 × g for 5 min, after which the pellet was resuspended in 0.5 mL of ice-cold PBS. The contents were thoroughly mixed by passing through a 200 µL pipette tip several times while keeping the tubes on ice throughout the process. In the next step, using a 200 µL tip, the cell suspension was transferred drop-wise into new 15 mL tubes containing 4.5 mL of ice-cold 70% ethanol, while continuously mixing the tubes on a vortex at low speed. The tubes were then incubated at 4 °C for 24 h to fix the cells. After fixation, the tubes containing the fixed cells were centrifuged for 5 min at 900 × g. The supernatant was carefully removed, and the cell pellet was resuspended in 5 mL of ice-cold PBS. Following another centrifugation for 5 min at 900 × g, the pellet was resuspended in 150 µL of PI Staining Solution (0.01 mg/mL PI, 0.1% Triton X-100, 0.01 mg/mL RNAse A), and the cells were incubated in the dark for 20 min. After the incubation period, the contents of the tubes were filtered through a nylon mesh into 1.5 mL tubes. The tubes were kept on ice and protected from light until analysis. Cells were analyzed using the Amnis® ImageStream®X Mk II IFC and the INSPIRE™ ImageStream®X System Software. Parameters were set to detect individual cells, targeting an optimal count of 15,000 cells, using a blue laser (excitation wavelength at 488 nm) with 5 mW power for the PI channel. Data analysis was conducted using the IDEAS® 6.2 Image Data Exploration and Analysis Software, employing histogram representation with a linearly scaled x-axis displaying the intensity of the fluorescent signal from PI. The y-axis represented the normalized frequency of cells exhibiting specific levels of fluorescent signal from PI. Results were presented as the percentage of cells in each phase of the cell cycle, specifically indicating the proportions of cells in the G_0_/G_1_, S, and G_2_/M phases.

**Monocyte-endothelial adhesion assay**

To assess the impact of short-term and long-term exposure of EA.hy926 cells to PFOA on monocyte adhesion to the endothelial cell monolayer, we used calcein AM-labeled human monocytic cell line U937, following the protocol established by Kokai et al. (2024) with minor modifications. For short-term exposure, EA.hy926 cells were plated in 35-mm dishes (0.7 × 10^6^ cells per dish) and exposed the following day to either vehicle (0.05% DMSO) or 1 μM, 10 μM, and 100 μM PFOA for 48 h. In long-term exposure experiments, EA.hy926 cells were harvested from each flask after 6 and 12 weeks of continuous exposure to either vehicle (0.05% DMSO) or 1 nM, 10 nM, and 100 nM PFOA, plated in 35-mm dishes (0.7 × 10^6^ cells per dish), and subjected to the corresponding treatments for an additional 48 h. A suspension of calcein AM-labeled U937 cells was added on top of the EA.hy926 monolayer (2 x 10^6^ cells per dish). After a 1-h incubation at 37 °C, the fluorescently labeled U937 cells bound to the EA.hy926 monolayer were photographed using a Zeiss Axio Imager.A2 fluorescence microscope with the appropriate filter (fluorescein isothiocyanate, FITC). Quantification was performed using NIH Image J software (<https://ImageJ.nih.gov/>), by counting the calcein AM-labeled U937 cells in the fluorescence images. The mean number of fluorescent cells obtained from 15 random digital images was calculated for each treatment and control, and the extent of U937 cell adhesion to the PFOA-exposed EA.hy926 monolayer was expressed relative to the control, which was set at 100%.

**Endothelial permeability assay**

Fibronectin-coated Transwell® units were used to assess the effect of short-term and long-term exposure of EA.hy926 cells to PFOA on endothelial permeability by measuring the passage of FITC-labeled dextran (FD40, Mw 40 kDa) through the EA.hy926 cell monolayer, according to the protocol described elsewhere (Kokai et al., 2020) with minor modifications. For short-term exposure, EA.hy926 cells were plated into fibronectin-coated inserts of the 24-well Transwell® plate (0.1 × 10^6^ cells per well; two wells per each treatment) and cultured in the complete growth medium for 6 days until a confluent semi-permeable barrier was formed, followed by exposure to either vehicle (0.05% DMSO) or 1 μM, 10 μM, and 100 μM PFOA for 48 h. In long-term exposure experiments, EA.hy926 cells were harvested from each flask after 5 and 11 weeks of continuous exposure to either vehicle (0.05% DMSO) or 1 nM, 10 nM, and 100 nM PFOA, plated into fibronectin-coated inserts of the 24-well Transwell® plate (0.1 × 10^6^ cells per well; two wells per each treatment) and subjected to the corresponding treatments for an additional 7 days to form a confluent semi-permeable barrier. FD40 in HBSS (1 mg/mL) was added to the upper compartment of the Transwell® plate, and the resulting fluorescence was measured in the lower compartment after 15, 30, 60, and 120 min using a Thermo Labsystems Fluoroskan Ascent fluorescence plate reader, with excitation at 485 nm and emission at 538 nm. Results were expressed as the average change in endothelial permeability over 5 min. The mean fluorescence for each treatment and control was calculated, with the permeability of PFOA-exposed EA.hy926 monolayer expressed as a percentage of change in FD40 fluorescence over 5 min in the lower compartment of the Transwell® plate relative to the control, which was set at 100%.

**Cell adhesion assay**

To investigate the effects of short-term and long-term PFOA exposure on the adhesion of EA.hy926 cells to the extracellular matrix, we conducted an adhesion assay using fibronectin-coated plates, following a modified protocol from Kokai et al. (2020). For short-term exposure, EA.hy926 cells were plated in 35-mm dishes (0.8 × 10^6^ cells per dish) and exposed to either vehicle (0.05% DMSO) or 1 μM, 10 μM, and 100 μM PFOA for 48 h. After exposure, cells were lifted into suspension using a 0.25% trypsin-EDTA solution for 4 min, followed by the addition of complete culture medium and centrifugation at 400 × g for 5 min. The pelleted cells were rinsed with PBS, centrifuged again, and resuspended in Adhesion Buffer (0.5% BSA, 1 mM CaCl_2_, 1 mM MgCl_2_, 0.2 mM MnCl_2_ in DMEM, pH 7.4). Cells in the resulting suspension were counted and plated in fibronectin-coated 12-well plates (0.4 × 10^6^ cells per well; two wells per each treatment) that had been blocked for 1 h with 1% BSA in PBS. In long-term exposure experiments, EA.hy926 cells were harvested from each flask after 6 and 12 weeks of continuous exposure to either vehicle (0.05% DMSO) or 1 nM, 10 nM, and 100 nM PFOA by trypsinization, followed by the addition of the complete culture medium and centrifugation at 400 × g for 5 min. Pelleted cells were rinsed with PBS, centrifuged again, resuspended in Adhesion Buffer, and plated in fibronectin-coated, BSA-blocked 12-well plates (0.4 × 10^6^ cells per well; one well per each flask). In both short-term and long-term experiments, plates were incubated for 30 min at 37 °C. Cells were then rinsed three times with DMEM and photographed using an Olympus inverted phase-contrast light microscope. The sulforhodamine B (SRB) assay was employed to quantify the number of attached cells (see below).

**ROS assay**

To assess the levels of reactive oxygen species (ROS) production following short-term and long-term exposure to PFOA, we used the cell-permeant ROS-sensitive probe 2',7'-dichlorodihydrofluorescein diacetate (H_2_DCFDA). This non-fluorescent probe becomes highly fluorescent 2',7'-dichlorofluorescein (DCF) upon cleavage of its acetate groups by intracellular esterases, which reduces its cellular permeability, and subsequent oxidation. For short-term exposure, EA.hy926 cells were plated in 96-well plates (2.5 × 10^6^ cells per well; 12 wells per treatment) and treated the following day with either vehicle (0.05% DMSO) or 1 μM, 10 μM, and 100 µM PFOA for 48 h. In long-term exposure experiments, EA.hy926 cells were harvested after 6 and 12 weeks of continuous exposure to either vehicle or 1 nM, 10 nM, and 100 nM PFOA. The harvested cells were then plated in 96-well plates (2.5 × 10^6^ cells per well; 12 wells per treatment) and exposed to the corresponding treatments for an additional 48 h. In both short-term and long-term experiments, cells were rinsed twice with HBSS, followed by the addition of 50 µM H_2_DCFDA in HBSS. The plates were incubated for 30 min in the dark at 37 °C, after which the cells were rinsed twice with PBS. PBS was then replaced with 10 mM HEPES buffer (pH 7.4) for 5 min. Fluorescence was measured using a Thermo Labsystems Fluoroskan Ascent fluorescence plate reader, with excitation at 485 nm and emission at 538 nm. The mean fluorescence for each treatment and control was calculated, and ROS production in PFOA-exposed EA.hy926 cells was expressed as a percentage of DCF fluorescence relative to the control, which was set at 100%. Subsequently, the results were normalized to the cell count in each well using the SRB assay.

**Sulforhodamine B assay**

The sulforhodamine B (SRB) assay was employed to quantify the number of adherent cells following the cell adhesion assay and the measurement of ROS levels, following the protocol established by Kokai et al. (2020).

**Cell migration assay**

To investigate the effect of short-term and long-term exposure to PFOA on endothelial cell migration, we performed a modified Boyden chamber assay, following the established protocol from our group (Stanic et al., 2024). For short-term exposure, EA.hy926 cells were plated in 35-mm dishes (0.6 × 10^6^ cells per dish) and exposed the following day to either vehicle (0.05% DMSO) or 1 μM, 10 μM, and 100 µM PFOA for 48 h. After exposure, cells were lifted into suspension using a 0.25% trypsin-EDTA solution for 4 min, followed by the addition of complete culture medium and centrifugation at 400 × g for 5 min. In long-term exposure experiments, EA.hy926 cells were harvested from each flask after 6 and 12 weeks of continuous exposure to either vehicle (0.05% DMSO) or 1 nM, 10 nM, and 100 nM PFOA by trypsinization, followed by the addition of the complete culture medium and centrifugation at 400 × g for 5 min. For both short-term and long-term exposure, the pelleted cells were resuspended in serum-free culture medium and counted. Cells were then plated on the top of polycarbonate membranes (pore size: 8 µm) of Transwell® inserts (5 × 10^4^ cells per insert in 100 µL). The subsequent steps were performed as previously described. Migration was quantified using a Zeiss Axio Imager.A2 fluorescence microscope (DAPI filter) by calculating the mean number of fluorescent cells from 10-15 randomly taken digital images per treatment group. The migration of PFOA-exposed EA.hy926 cells was expressed as the percentage of fluorescently stained cells on the polycarbonate membrane of the Transwell® plate relative to the control, which was set at 100%.

**Tube formation assay**

The tube formation assay was conducted to evaluate the effect of both short-term and long-term exposure of EA.hy926 cells to PFOA on *in vitro* angiogenesis, following the protocols outlined in our previous publications (Kokai et al., 2022; Stanic et al., 2024) with minor modifications. For short-term exposure, EA.hy926 cells were plated in 35-mm dishes (0.6 × 10^6^ cells per dish) and exposed to either vehicle (0.05% DMSO) or 1 μM, 10 μM, and 100 µM PFOA for 48 h. After exposure, cells were lifted into suspension using a 0.25% trypsin-EDTA solution for 4 min. The suspension was then supplemented with complete culture medium, followed by centrifugation at 400 × g for 5 min, a quick rinse with PBS, and another centrifugation at 400 × g for 5 min. In long-term exposure experiments, cells were harvested from each flask after 6 and 12 weeks of continuous exposure to either vehicle (0.05% DMSO) or 1 nM, 10 nM, and 100 nM PFOA. The harvesting process involved trypsinization, followed by the addition of complete culture medium and centrifugation at 400 × g for 5 min, a quick rinse with PBS, and a final centrifugation at 400 × g for 5 min. In both short-term and long-term exposure experiments, the pelleted cells were resuspended in serum-free culture medium and counted. Corresponding treatments were added to the appropriate number of cells (0.5 × 10^6^ cells per mL). From this point on, the assay proceeded as described earlier. Morphometric analysis of capillary-like tubes was performed using the Angiogenesis Analyzer, an Image J toolset (Carpentier et al., 2020), by analyzing three photographic fields per well. The extent of angiogenesis in PFOA-exposed EA.hy926 cells was quantified based on the total length of tubes, number of nodes, number of junctions, and total branching length, expressed relative to the control, which was set at 100%.

**Statistical analysis**

Statistical analysis was conducted using the Prism 8 software package (GraphPad Software, Inc., La Jolla, CA, USA) by one-way analysis of variance (ANOVA) with Dunnett’s multiple comparisons post-hoc test. A *p*-value of <0.05 was considered significant.

**SUPPLEMENTARY FIGURES**

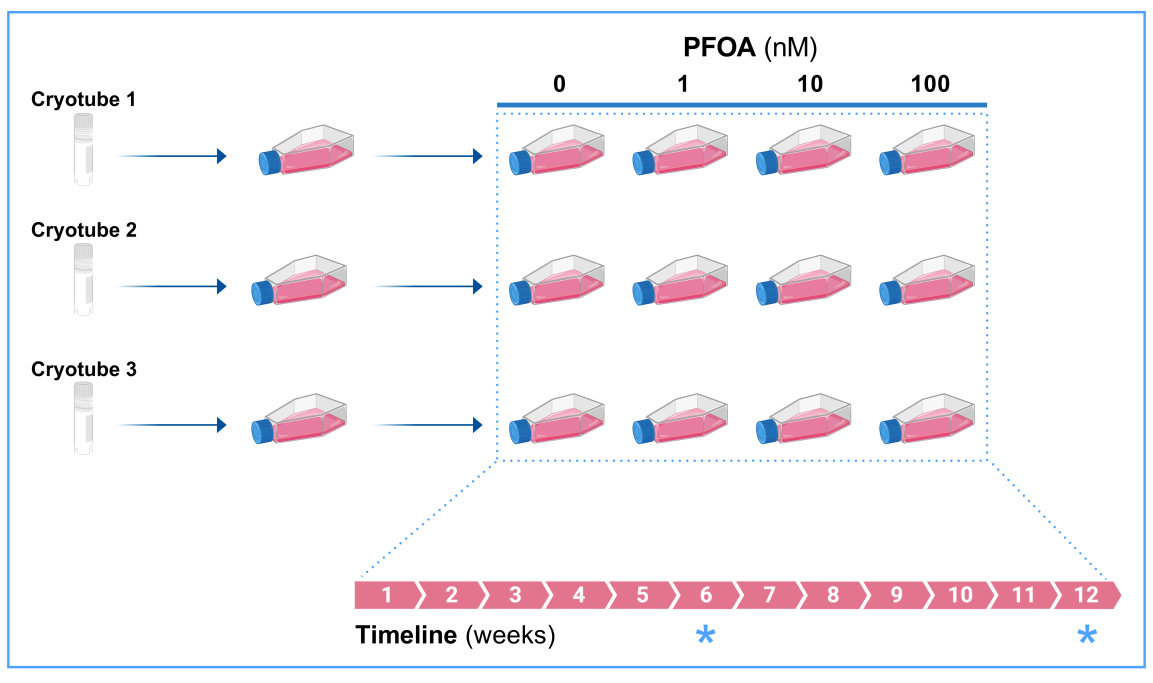

**Supplementary Figure 1.** Schematic representation of the 12-week-long exposure of EA.hy926 cells to 1 nM, 10 nM, and 100 nM PFOA. Three different cryopreserved stock vials of EA.hy926 cells (biological replicates) were thawed and cultured in separate 25 cm^2^ cell culture flasks for two weeks. After this period, cells from each flask were divided into four flasks, resulting in a total of 12 flasks. Over the subsequent 12 weeks, EA.hy926 cells were subcultured twice a week, on Tuesdays and Fridays, and exposed to either vehicle (0.05% DMSO – control group) or three nanomolar concentrations of PFOA (1, 10, and 100 nM in 0.05% DMSO – treatment groups) 3 h after seeding to minimize the effects of treatments on cell attachment to the flask surface. After 6 and 12 weeks of continuous exposure (indicated by the blue asterisk), cells were collected directly from the flasks and used for various apical assays and mRNA sequencing or plated in appropriate cell culture plates for other apical assays, as indicated. Image created with BioRender.

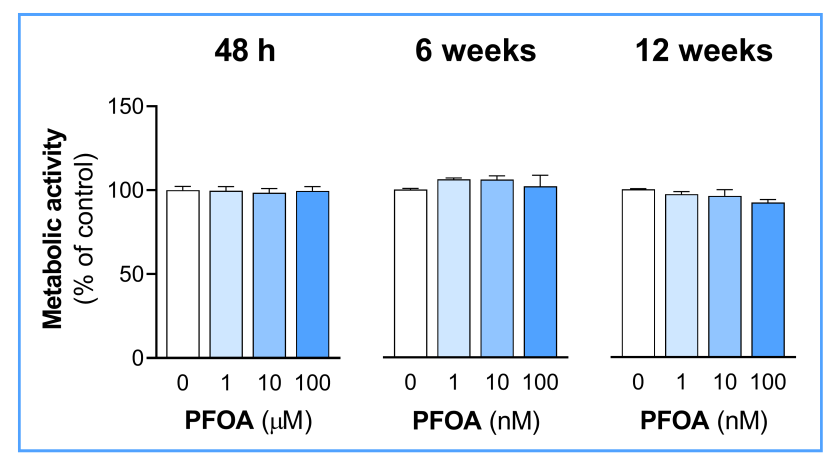

**Supplementary Figure 2.** Effect of short-term and long-term exposure of EA.hy926 cells to PFOA on metabolic activity. EA.hy926 cells were exposed to 1 μM, 10 μM, and 100 μM PFOA for 48 h (short-term exposure) and 1 nM, 10 nM, and 100 nM PFOA for 6 and 12 weeks (long-term exposure). Metabolic activity was assessed using the alamarBlue™ assay. Results were expressed relative to the vehicle-treated control (100%). Each data bar represents the mean ± SEM of 4 independent experiments (short-term exposure) or 3 cell culture flasks (long-term exposure).

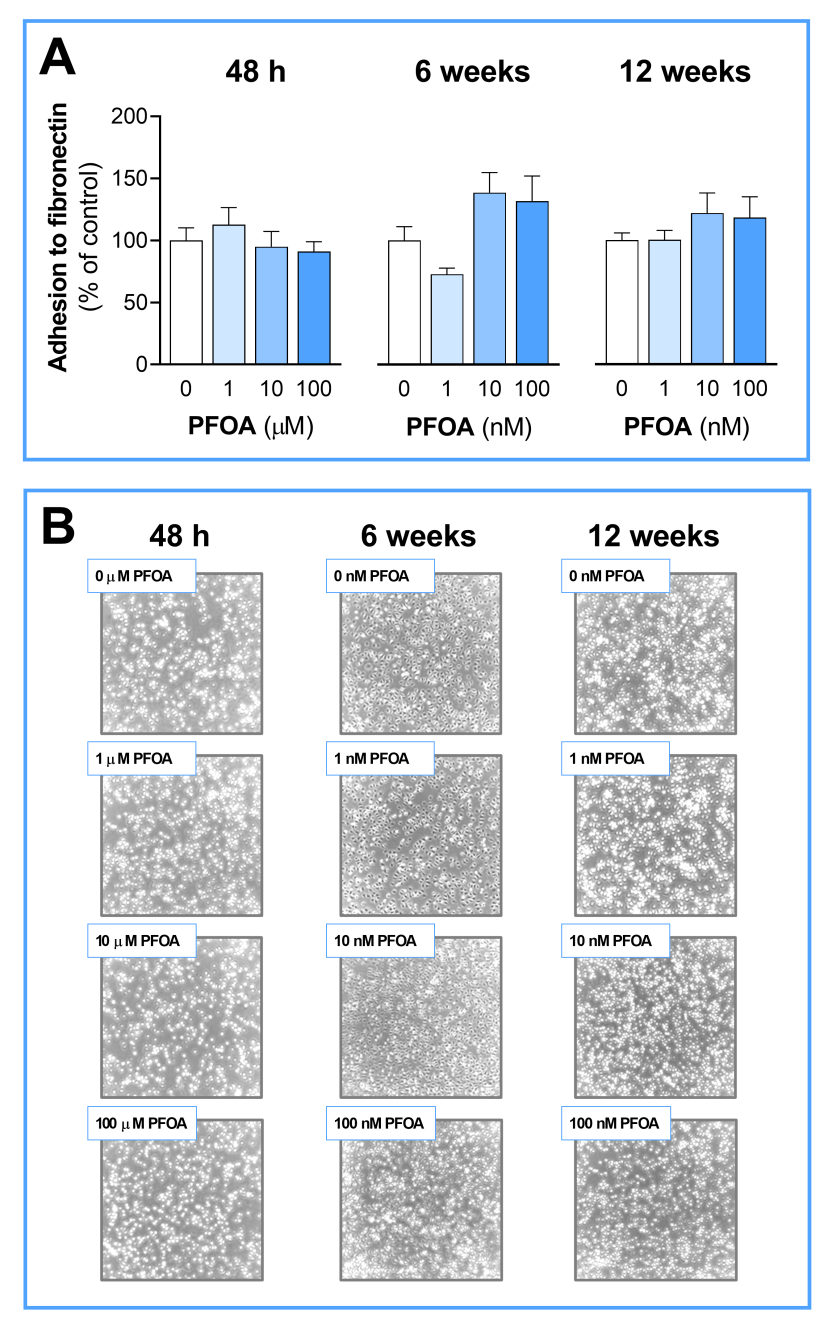

**Supplementary Figure 3.** Effect of short-term and long-term exposure of EA.hy926 cells to PFOA on cell adhesion to fibronectin. EA.hy926 cells were exposed to 1 μM, 10 μM, and 100 μM PFOA for 48 h (short-term exposure) and 1 nM, 10 nM, and 100 nM PFOA for 6 and 12 weeks (long-term exposure). The extent of cell adhesion to fibronectin was assessed after 30 min using an inverted light microscope, and the number of adhered cells was quantified with the SRB assay. **(A)** Results were expressed relative to the vehicle-treated control (100%). Each data bar represents the mean ± SEM of 4 independent experiments (short-term exposure) or 3 cell culture flasks (long-term exposure). **(B)** Representative photomicrographs are shown.

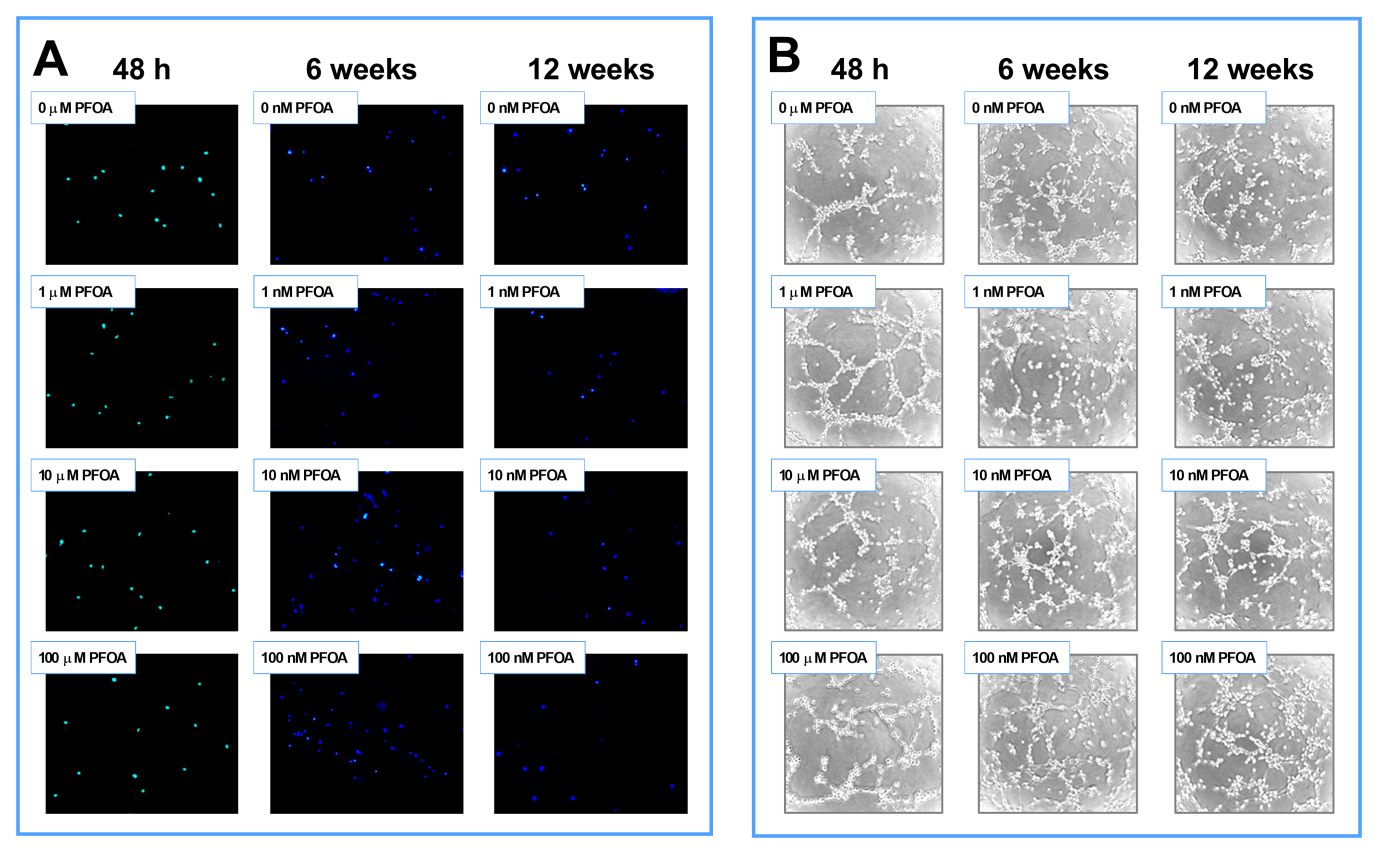

**Supplementary Figure 4.** Effect of short-term and long-term exposure of EA.hy926 cells to PFOA on cell migration and endothelial tube formation. EA.hy926 cells were exposed to 1 μM, 10 μM, and 100 μM PFOA for 48 h (short-term exposure) and 1 nM, 10 nM, and 100 nM PFOA for 6 and 12 weeks (long-term exposure). **(A)** The extent of cell migration was assessed using the modified Boyden chamber assay. Cells that migrated through the polycarbonate membrane of the inner compartment of the Transwell® plate after 5 h were stained with the fluorescent dye DAPI and observed under a fluorescence microscope. Representative fluorescent photomicrographs are shown. The number of migrated cells was quantified using Image J. **(B)** Endothelial tube formation in a three-dimensional matrix (growth factor-reduced Cultrex BME) was observed under an inverted light microscope after 8 h. Representative photomicrographs are shown. The obtained digital images were analyzed using the Angiogenesis Analyzer plug-in in Image J.

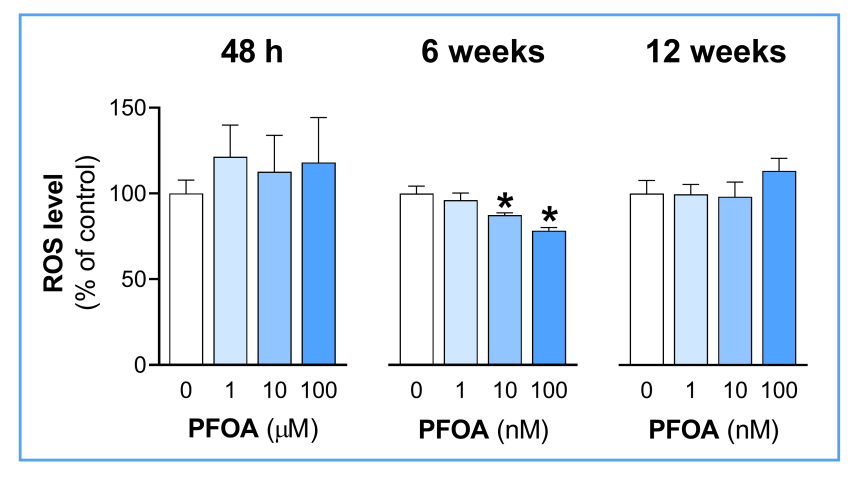

**Supplementary Figure 5.** Effect of short-term and long-term exposure of EA.hy926 cells to PFOA on ROS levels. EA.hy926 cells were exposed to 1 μM, 10 μM, and 100 μM PFOA for 48 h (short-term exposure) and 1 nM, 10 nM, and 100 nM PFOA for 6 and 12 weeks (long-term exposure). ROS production was assessed using H_2_DCFDA. Results were expressed relative to the vehicle-treated control (100%). Each data bar represents the mean ± SEM of 4 independent experiments (short-term exposure) or 3 cell culture flasks (long-term exposure). **p* <0.05 *vs.* control.

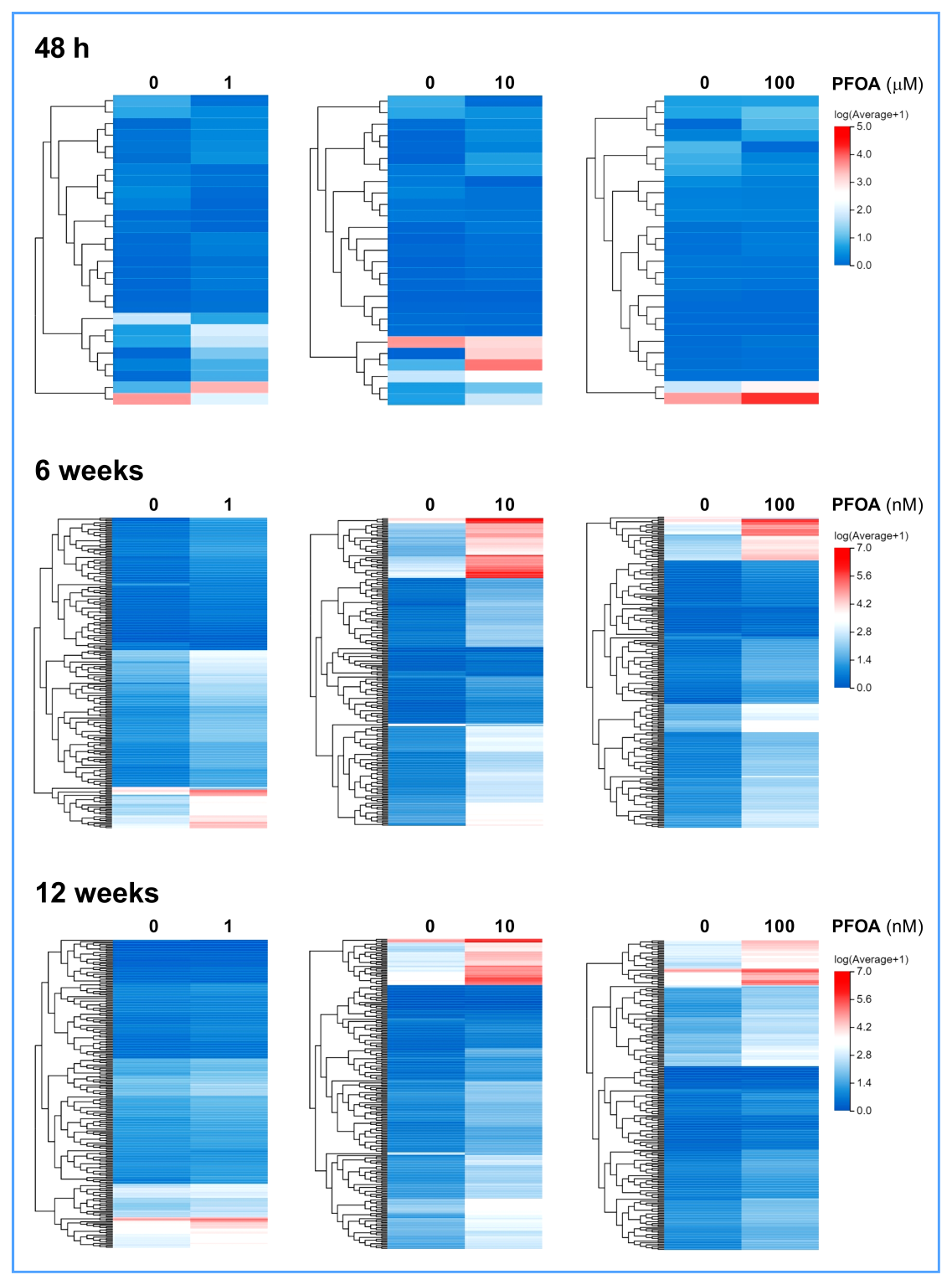

**Supplementary Figure 6**. Heatmaps illustrating DEGs following short-term (48 hours) and long-term (6 and 12 weeks) exposure of EA.hy926 cells to specified concentrations of PFOA. The cluster analysis highlights transcripts that exhibited significant changes at each concentration. Upregulated genes are represented in shades of red, downregulated genes in shades of blue, and genes with no change in expression are shown in white.

**SUPPLEMETARY REFERENCES**

Carpentier G, Berndt S, Ferratge S, Rasband W, Cuendet M, Uzan G, Albanese P. 2020. Angiogenesis Analyzer for ImageJ – A comparative morphometric analysis of “Endothelial Tube Formation Assay” and “Fibrin Bead Assay.” Scientific Reports, 10(1):11568. doi: 10.1038/S41598-020-67289-8

Kokai D, Stanic B, Samardzija Nenadov D, Pogrmic-Majkic K, Tesic B, Fa S, Andric N. 2020. Biological effects of chronic and acute exposure of human endothelial cell line EA.hy926 to bisphenol A: New tricks from an old dog. Chemosphere, 256:127159. doi: 10.1016/j.chemosphere.2020.127159

Kokai D, Stanic B, Tesic B, Samardzija Nenadov D, Pogrmic-Majkic K, Fa Nedeljkovic S, Andric N. 2022. Dibutyl phthalate promotes angiogenesis in EA.hy926 cells through estrogen receptor-dependent activation of ERK1/2, PI3K-Akt, and NO signaling pathways. Chemico-Biological Interactions, 366:110174. doi: 10.1016/j.cbi.2022.110174

Kokai D, Markovic Filipovic J, Opacic M, Ivelja I, Banjac V, Stanic B, Andric N. 2024. In vitro and in vivo exposure of endothelial cells to dibutyl phthalate promotes monocyte adhesion. Food and Chemical Toxicology, 188:114663. doi: 10.1016/j.fct.2024.114663

Stanic B, Kokai D, Markovic Filipovic J, Tomanic T, Vukcevic J, Stojkov V, Andric N. 2024. Vascular endothelial effects of dibutyl phthalate: In vitro and in vivo evidence. Chemico-Biological Interactions, 399:111120. doi: 10.1016/j.cbi.2024.111120
